# Vascular burden is associated with a decline in default-mode and global resting-state functional connectivity in individuals at risk for Alzheimer’s disease

**DOI:** 10.1101/2020.04.10.036202

**Authors:** Theresa Köbe, Alexa Pichet Binette, Jacob W. Vogel, Pierre-François Meyer, John C. S. Breitner, Judes Poirier, Sylvia Villeneuve, for the Presymptomatic Evaluation of Novel or Experimental Treatments for Alzheimer Disease (PREVENT-AD) Research Group

## Abstract

**Introduction:** Cross-sectional studies suggest that cardiovascular risk factors and Alzheimer’s disease (AD) biomarkers are associated with abnormal brain resting-state functional connectivity in aging and AD; however, evidence is missing regarding longitudinal changes in functional connectivity. In this study, we investigate whether cholesterol levels and blood pressure are associated with changes in functional connectivity over time in asymptomatic individuals at risk for AD. The analyses were repeated with cerebral β-amyloid (Aβ) and *tau* deposition in a subset of the participants.

**Methods:** The study sample included 247 cognitively unimpaired individuals (185 women/ 62 men; mean [SD] age of 63 [5.3] years) of the PREVENT-AD cohort with a parental or multiple-sibling history of sporadic AD. Plasma total-, HDL-, and LDL-cholesterol and systolic and diastolic blood pressure were measured at baseline. Global brain functional connectivity, and connectivity from canonical functional networks, were computed from resting-state functional MRI obtained at baseline and up to four years of annual follow-ups, using a predefined functional parcellation. A subset of participants underwent *tau*-PET ([^18^F]Flortaucipir) and Aβ-PET ([^18^F]NAV4694). Vascular and AD measures were examined as predictors of brain functional connectivity changes in linear mixed-effects models.

**Results:** Higher total-cholesterol and LDL-cholesterol levels were associated with greater reduction of functional connectivity in the default-mode network over time. In addition, while overall whole-brain functional connectivity showed an increase over time across the entire sample higher diastolic blood pressure was associated with reduction in whole-brain functional connectivity. The associations were similar when the analyses were repeated using two other functional brain parcellations. The findings with total-cholesterol and diastolic blood pressure were also similar but attenuated when performed in a subsample of participants with PET (n=91), whereas AD biomarkers were not associated with changes in functional connectivity over time in this subsample.

**Conclusion:** These findings provide evidence that vascular burden is associated with a decrease in brain functional connectivity over time in older adults with elevated risk for AD. The impact of vascular risk factors on functional brain changes might precede AD pathology-related changes.

## 1 Introduction

Alzheimer’s disease (AD) is a multifactorial disease that is characterized not only by pathological protein aggregation in the brain (Aβ and tau), but also by early vascular dysfunctions and changes in brain functional connectivity.^1-3^ Brain regions that show a temporal correlation in blood-oxygenation-level-dependent (BOLD) signals, measured with resting-state functional magnetic resonance imaging (rs-fMRI), have been proposed to be intrinsically functionally connected. Consistent resting-state functional connectivity (RSFC) networks have been identified in a large number of studies, using different brain parcellations.^4-6^ Both aging^7^ and neurodegenerative disease^8^ are characterized by alterations within these networks. With age, RSFC appears to decline within most networks but tends to increase between networks.^7^ It has in fact recently been suggest that global and default-mode network (DMN) RSFC tend to increase up to the seventh decade, followed thereafter by an accelerated age-related decline.^9^ Certain alterations in RSFC have been consistently linked to mild cognitive impairment and AD dementia, particularly with lower RSFC in the DMN, but also in the salience (SAL) and limbic (LIM) networks, as well as changes in global RSFC.^10-16^ AD-related changes in RSFC already appear in the asymptomatic stage, years before disease onset.^17,18^ Cognitively normal individuals with a family history of sporadic AD and individuals with subjective cognitive decline also present RSFC alterations.^19-21^ Functional changes associated with AD risk are not only linked to a reduction of RSFC, but also to compensatory increase in RSFC.^17,20^

Vascular risk factors (VRF) such as dyslipidemia and hypertension are thought to impair healthy aging and to increase AD risk.^22,23^ Cross-sectional studies have demonstrated that higher vascular burden is associated with reduced cerebral metabolism, reduced cerebrovascular reactivity and an associated reduction in functional connectivity.^24-27^ Hypertension has been suggested to impair RSFC in older adults.^28^ Higher total-cholesterol levels have been associated with both lower and higher RSFC within the DMN and lower RSFC within the SAL network,^29,30^ but associations have not been found in all studies.^26^ VRFs might therefore reduce brain functional health, which could increase vulnerability to AD pathology and/or cognitive impairments. VRFs being present prior to AD pathology in most individuals, the impact of VRF on brain integrity could precede the one related to AD pathology.^22^

In the current study, we aimed to investigate how RSFC alterations develop over time in the presence of VRFs and AD biomarkers. We applied longitudinal rs-fMRI to delineate related changes in functional connectivity in cognitively normal individuals, who had a 2-3 fold higher risk for AD owing to a first-degree family history of AD.^31^ Our main hypothesis was that individuals with higher vascular burden will show a decrease in RSFC over time, and that this reduction would be predominant in the DMN. Second, we hypothesized that higher Aβ and *tau* burden will result in similar effects, with *tau* also being associated with change in the limbic network.

## 2 Materials and methods

### Participants and Study design

Study participants were recruited within the *PResymptomatic EValuation of Experimental or Novel Treatments for Alzheimer Disease* (PREVENT-AD) study, an ongoing longitudinal observational study initially enrolling 385 individuals.^32^ Most of this dataset is now openly accessible (https://openpreventad.loris.ca/). Study participants met the following inclusion criteria: (a) parental or multiple-sibling history of AD-like dementia, (b) ≥60 years old at enrollment or 55-59 years old if less than 15 years away from youngest affected relative’s age of symptom onset, (c) no major neurological diseases and (d) normal cognition. All participants underwent neuropsychological testing at baseline using the Repeatable Battery for the Assessment of Neuropsychological Status (RBANS),^33^ Clinical Dementia Rating (CDR)^34^ and the Montreal Cognitive Assessment (MOCA)^35^ to assess normal cognition. In the few cases of ambiguous CDR or MOCA assessments (n=15), participants were further evaluated with a more extensive neuropsychological test battery that was reviewed by neuropsychologists (including SV) and physicians (including JCSB) to ensure normal cognition. **Figure 1** provides an overview of the study.

**Figure 1:**
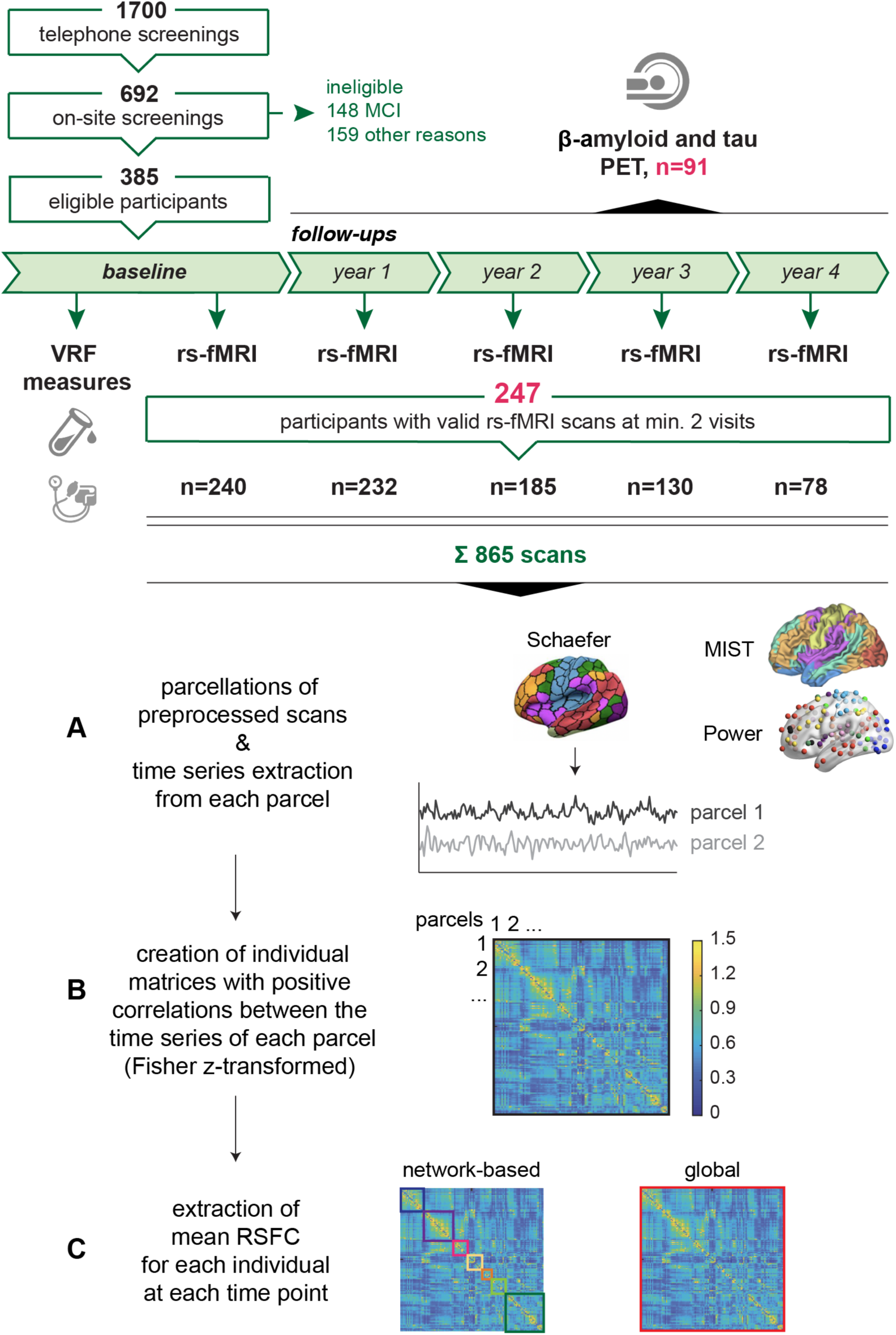
Study flow chart and fMRI analyses. In total, 385 eligible PREVENT-AD participants were followed annually over up to 4 years. Vascular risk factors were assessed at baseline. Resting-state functional magnetic resonance images (rs-fMRI) were acquired at baseline and follow-up year 1 to 4. Finally, 247 participants with 865 scans were included in the current study, who had at least 2 valid rs-fMRI scans across baseline (n=240 scans), follow-up year 1 (n=232), year 2 (n=185), year 3 (n=130) and year 4 (n=78). For 7 participants the first valid MRI scan was available at follow-up year 1 (n=5) and 2 (n=2) instead of at baseline. (A) Individual time series of preprocessed scans were extracted from each parcel, defined by the Schaefer parcellations. Analyses were repeated with the MIST and Power atlas. (B) Correlation matrices were obtained by correlating the time series of each parcel with one another. Correlation matrices were Fisher z-transformed and thresholded by only keeping positive correlations. (C) Extracted mean correlations represent the indirect measure of network-based (multicolored squares) and global (red square) resting-state functional connectivity (RSFC). A subsample of 91 participants that had two rs-fMRI scans underwent Aβ and tau PET scanning during the course of the study.

In the current longitudinal analyses, 247 participants were included who had at least 2 valid fMRI scans.

### Standard protocol approvals, registrations, and patient consents

The study was approved by the McGill University Faculty of Medicine Institutional Review Board. All participants received detailed study instructions and gave written consent prior to participation.

### VRF assessment

All participants were examined medically, venous blood samples (non-fasting) and blood pressure were taken at baseline (see Supplementary material for more details). Plasma levels of total-, high-density (HDL)- and low-density lipoprotein (LDL)-cholesterol were measured by standard enzymatic methods (CHOD-PAP; Beckman Coulter, Synchron LX®, UniCel® DxC 600/800 System and Synchron® Systems Lipid Calibrator). Blood pressure was assessed while seated in a standardized procedure using an automatic sphygmomanometer (Connex® ProBP™ 3400; Welch Allyn).

### APOE Genotyping

Genomic DNA was extracted from whole blood and *APOE* genotype was determined using the PyroMark Q96 pyrosequencer (Qiagen, Toronto, ON, Canada), as described previously.^36^ Participants were classified as *APOE* ε4 carriers (one or two ε4 alleles) or noncarriers.

### MRI acquisition

Participants underwent MRI annually (at baseline and at follow-ups from 1 to 4 years) on a 3T Siemens Trio scanner at the Brain Imaging Centre of the Douglas Mental Health University Institute (Montreal, Canada). Structural T1-weighted images were obtained with the following parameters: TR=2300ms, TE=2.98ms, number of slices=176, slice thickness=1mm. For resting-state functional MRI (rs-fMRI) scans, two consecutive functional T2*-weighted images (each “run” with 150 volumes lasting 5min and 45s) were acquired using a blood-oxygen-level-dependent (BOLD) sensitive, single-shot echo planar sequence (TR=2000ms; volumes=150; TE=30ms; FA=90°; matrix size=64×64; voxel size=4x 4×4mm3; 32 slices). The participants were asked to keep their eyes closed and to remain as still as possible during scanning.

### MRI analyses

Preprocessing and analyses of the rs-fMRI data has been described previously,^20^ using the Neuroimaging Analysis Kit (http://niak.simexp-lab.org, v0.12.17), GNU Octave (v4.0), and the Minc toolkit (http://www.bic.mni.mcgill.ca/ServicesSoftware/ServicesSoftwareMinc ToolKit, v0.3.18). Briefly, the preprocessing of functional scans comprised motion-correction, slice-time correction, temporal filtering (0.01Hz high-pass cut-off), non-linear spatially normalization to the Montreal Neurological Institute ICBM152 symmetric template, resampling to 2mm^3^, and spatial smoothing with a 6mm full-width-half-maximum (FWHM) Gaussian kernel. Noise due to motion, slow time drifts, and average signals in white matter and the lateral ventricles was removed from the functional signal by multiple regression (http://niak.simexplab.org/pipe_preprocessing.html).^20,37^ In addition, time frames with in-scanner head motion (mean framewise displacement (FD)) above 0.5mm were removed (scrubbed) from individual time series, along with one adjacent frame prior and two following frames after, to minimize artefacts caused by excessive motion.^38^ Further descriptions of the pipeline can be found on the NIAK website (http://niak.simexp-lab.org/build/html/PREPROCESSING.html). Moreover, rs-fMRI images passed quality control if: (a) at least one out of two functional runs remained with a minimum of 70 frames (140s) after scrubbing; (b) no image artefacts were found during visual examination; (c) they were correctly co-registered to structural MRI and ICBM152 template (spatial correlation r>0.75, NIAK preprocessing report).

In total, 1152 rs-fMRI scans were acquired within the PREVENT-AD cohort at 5 acquisition time points (baseline and follow-ups at year 1 to 4), of which 198 scans failed quality control, leaving 954 valid scans. For the current longitudinal analyses, participants were required to have at least 2 valid scans at different time points, resulting in 865 scans from 247 individuals (median [interquartile range] number of scans, 4 [2-4] and follow-up time, 3 [2-4] years) (see **Table 1**).

**Table 1:** Overview MRI flow

| Numbers of: | BL | FU year 1 | FU year 2 | FU year 3 | FU year 4 | Total |
| --- | --- | --- | --- | --- | --- | --- |
| **acquired MRI scans** | 363 | 294 | 232 | 169 | 94 | 1152 |
| **failed quality control (reason<sup>a/b/c</sup>)** | -51<br>(50<sup>a</sup>/1<sup>b</sup>) | -51<br>(48<sup>a</sup>/3<sup>c</sup>) | -43<br>(42<sup>a</sup>/1<sup>c</sup>) | -37<br>(37<sup>a</sup>) | -16<br>(16<sup>a</sup>) | -198<br>(193<sup>a</sup>/1<sup>b</sup>/4<sup>c</sup>) |
| **valid MRI scans** | 312 | 243 | 189 | 132 | 78 | 954 |
| **individuals with at least 2 valid MRI scans (with one/ two runs)** | 240<br>(20/220) | 232<br>(17/215) | 185<br>(28/157) | 130<br>(13/117) | 78<br>(12/66) | **865**<br>**(90/775)**<br>**= 247 unique individuals** |

Reasons for failure of quality control: <sup>a</sup> <70 unscrubbed frames per run; <sup>b</sup> image artefacts; <sup>c</sup> registration failure. For 7 participants the first valid MRI scans were available at follow-up year 1 (n=5) and year 2 (n=2) instead at baseline. BL, baseline; FU, follow-up.

For *a priori* network-based and global connectivity analyses we used the “Schaefer parcellation”.^4^ This parcellation scheme includes 400 predefined regions-of-interest (ROIs) classified into 7 neocortical functional networks (default mode [DMN], salience and ventral attention [SAL/VAN], fronto-parietal [FPN], limbic [LIM], dorsal attention [DAN], visual [VIS] and somatomotor network [SM]). To ensure the robustness of our findings, we repeated our analyses, using 2 different predefined parcellation schemes. The second parcellation was the “MIST parcellation”6, consisting of 444 symmetric ROIs that belong to 7 functional networks (DMN, SAL/VAN, FPN, LIM, VIS, SM and cerebellum). We restricted the analyses to neocortical functional networks to match the Schaefer parcellation and therefore excluded the cerebellum. The third atlas was the “Power parcellation”, including 264 ROIs, made up of 5mm radius spheres^39^ set around network coordinates predefined by Power et al.^5^; thus, varying in shape and number from the other two parcellation schemes. Here, we restricted our network analyses to the 7 matching neocortical functional networks, i.e. DMN, SAL, VAN, FPN, DAN, VIS and SM, and included 8 additional spherical ROIs (5mm radius) encompassing the LIM as done previously.^39^

For each atlas, the average time series from each parcel was extracted, correlated with one another (Pearson) and Fisher z-transformed, using Matlab (R2018b). Hence, single-subject correlation matrices were created (400×400, 444×444 and 272×272 for the Schaefer, MIST and Power parcellation, respectively). For participants who had 2 valid rs-fMRI runs per visit, correlation matrices were averaged after processing. Results were essentially unchanged when correcting our subsequent statistical models for the number of valid runs (1 or 2) at each visit. Given the ambiguous interpretation of negative correlations, correlation matrices were thresholded to keep only positive correlations.^40^ Finally, correlation estimates from parcels of the same network were averaged, providing a mean RSFC within each functional network. In addition, we computed the global (whole-brain) RSFC for each participant, corresponding to the average connectivity of each ROI to the entire rest of the cerebrum (Schaefer: 400×400; MIST: 387×387 (cerebellum ROIs excluded); and Power: 232×232 (cerebellum and as uncertain defined ROIs excluded)). We investigated both network-specific and global functional connectivity given that specific and global brain connections seem to be differentially associated to brain alterations and AD-related biomarkers.^12^ An overview of the rs-fMRI processing and analyses is represented in **Figure 1**.

### PET assessment and processing

In a subsample of 91 (37%) participants, PET scans using [^18^F]NAV4694 (NAV) for amyloid-β and [^18^F]AV1451 (Flortaucipir) for *tau* were acquired at the McConnell Brain Imaging Centre of the Montreal Neurological Institute (MNI) on a PET Siemens/CTI high-resolution scanner during the course of the study.^23^ Static acquisition frames were obtained for Aβ at 40-70 min. and for *tau* at 80-100 min. post-injection. More information about PET assessment are available in the supplement. PET data were pre-processed using a standard pipeline (see https://github.com/villeneuvelab/vlpp for details). Briefly, 4D PET images were averaged and linearly co-registered to individual’s T1-weighted images, before being masked to exclude CSF binding and smoothed with a 6mm^3^ Gaussian kernel. Individual T1-weighted images were segmented based on the Desikan-Killiany atlas using the semiautomated FreeSurfer processing stream version 5.3.^41^ Standardized uptake value ratios (SUVR) were computed for Aβ^42^ and *tau*^43^ by dividing the tracer uptake by cerebellar gray matter and inferior cerebellar gray matter uptake, respectively. We restricted the ROI analyses to FreeSurfer-derived AD-typical regions, i.e. weighted mean SUVRs from frontal, temporal, parietal and posterior cingulate cortex for a global Aβ quantification^42^ and from the entorhinal cortex for *tau* quantification.^44^

### Statistical analyses

Analyses were performed with R 3.5.2 (The R Foundation) and SPSS 24.0 (IBM Corp., Armonk, NY). Two-tailed *p*-values <0.05 were considered to be significant. First, we run cross-sectional general linear models, corrected for baseline age, sex, vascular medication (intake or non-intake of drugs against dyslipidemia and/or hypertension) and mean FD, to test for associations between VRFs and network/global RSFC at baseline. In our principal analyses, we run linear mixed-effects models (lmer function of the *lme4* package^45^) to test whether VRFs at baseline were associated with longitudinal changes in RSFC, separately in each network and globally throughout the whole cerebrum. All models were corrected for baseline age, sex and vascular medication and mean FD at each visit as well as their interactions with time. Follow-up time was operationalized individually as years from baseline. We included all VRFs in one model, except for total- and LDL-cholesterol, which showed high multicollinearity (variance inflation factor > 5: total-cholesterol ∼9 and LDL-cholesterol ∼8). Hence, we performed 2 separate models for each RSFC network and global connectivity, using the following equations:

**Model A)** RSFC ∼ total-cholesterol * time + HDL-cholesterol * time + systolic blood pressure *time + diastolic blood pressure * time + covariates * time + (time|ID) **Model B)** RSFC ∼ LDL-cholesterol * time + HDL-cholesterol * time + systolic blood pressure *time + diastolic blood pressure * time + covariates * time + (time|ID)

Results from each parcellation (and for each Model A and B if applicable) were separately evaluated for significance at a False Discovery Rate (FDR) corrected *p*-values <0.05. When results survived FDR correction it was explicitly stated in the manuscript.

To test for an overall change in RSFC over time, without taking VRFs into account, we also considered linear mixed-effects models including time as the independent variable, adjusted for age, sex and mean FD.

In exploratory analyses, we tested for an association of Aβ and *tau* deposition with changes in RSFC in a subgroup of PET participants (n=91), using linear mixed-effects models corrected for age, sex and mean FD. Here, the RSFC time variable was centered on the PET visit for each individual, since the PET scans were acquired during the course of the study. Analyses with main VRF findings on RSFC were repeated in the same subsample of PET participants to test if these associations are comparable with PET findings on RSFC changes (here *p*-values <0.1 were reported as trend-level).

In all linear mixed-effects models, we included individual RSFC intercepts and slopes as random effects. All continuous variables were z-transformed prior to model estimation. All models used the restricted maximum likelihood method and were fit with an unstructured variance-covariance and type III sum of squares. Denominator degrees of freedom were calculated with the Satterthwaite approximations.

## 3 Results

### Participant characteristics

In total, 247 participants were included in the current longitudinal analyses. At study entry, they were on average 63 years old, 75% were female, they had a mean education of 15.4 years and 39% carried at least one *APOE* ε4 allele. Participants were on average 11 years away from their relative’s age at AD dementia symptom onset. The subgroup of 91 participants with PET showed low to moderate levels of Aβ and *tau* deposition. See **Table 2** for further details. Participants were generally in good health, but nevertheless presented a wide range of VRFs. Abnormal levels of VRFs were predominantly recorded for total-cholesterol levels (>5.2 mmol/L; 63% of the participants), LDL-cholesterol (>3.4 mmol/L; 38%), systolic blood pressure (≥130 mmHg; 43%), diastolic blood pressure (≥80 mmHg; 29%) and body mass index (>30 kg/m^2^; 14%). About 4% of the study sample was diagnosed with diabetes and 4% of the participants reported current smoking.

**Table 2:** Participant characteristics at baseline

| Characteristics | Mean (SD; range) |  |  |  |  |
| --- | --- | --- | --- | --- | --- |
| Sample size [No.] | 247 |  |  |  |  |
| Age [years] | 62.90 (5.31; 55–83) |  |  |  |  |
| Sex [No. women (%)] | 185 (75) |  |  |  |  |
| Education [years] | 15.37 (3.58; 7–29) |  |  |  |  |
| APOE ε4 [No. carrier, ≥1 ε4 allele (%)] | 96 (39) |  |  |  |  |
| spEYO [years]<sup>a</sup> | -10.93 (7.58; -28–16) |  |  |  |  |
| Plasma lipids [mmol/L] |  |  |  |  |  |
| Total-cholesterol | 5.48 (0.99; 2.20–8.30) |  |  |  |  |
| HDL-cholesterol | 1.57 (0.40; 0.74–3.04) |  |  |  |  |
| LDL-cholesterol<sup>b</sup> | 3.09 (0.85; 0.78–5.77) |  |  |  |  |
| Arterial blood pressure [mm Hg] |  |  |  |  |  |
| Systolic blood pressure | 127.50 (15.87; 82–174) |  |  |  |  |
| Diastolic blood pressure | 73.46 (9.17; 51–110) |  |  |  |  |
| Medication [No. (%)] | 80 (32.4) |  |  |  |  |
| Lipid-lowering drugs | 24 (9.7) |  |  |  |  |
| Antihypertensive drugs | 32 (13.0) |  |  |  |  |
| Lipid-lowering & antihypertensive drugs | 24 (9.7) |  |  |  |  |
| AD PET biomarkers, median (IQR), [range]<sup>c</sup> |  |  |  |  |  |
| Aβ, [<sup>18</sup>F]NAV-4694 SUVR | 1.21 (1.14-1.29), [1.05-2.81] |  |  |  |  |
| tau, [<sup>18</sup>F]AV-1451 SUVR | 1.04 (0.98-1.14), [0.84-1.66] |  |  |  |  |
| Mean FD<sup>d</sup><br>(BL, FU1, FU2, FU3, FU4) | 0.22 (0.05; 0.09–0.33) | 0.22 (0.05; 0.09–0.35) | 0.23 (0.05; 0.12–0.34) | 0.23 (0.04; 0.10–0.33) | 0.23 (0.05; 0.11–0.34) |

<sup>a</sup>The spEYO could not be calculated for 16 participants, due to missing information about the family's age of onset. <sup>b</sup>LDL-cholesterol values were missing for 7 participants, due to laboratory measurement failures. <sup>c</sup>A subsample of 91 participants underwent β-amyloid (Aβ ) and tau PET scanning during the course of the study. <sup>d</sup>The sample size of the fMRI data differs between BL (n=240), FU year 1 (n=232), FU year 2 (n=185); FU year 3 (n=130); FU year 4 (n=78). APOE ε4, apolipoprotein E ε4; BL, baseline; FD, frame-wise displacement; FU, follow-up; EYO, estimated years to symptom onset of sporadic Alzheimer's disease, calculated by subtracting the age at family's symptom onset from the participant's age at baseline.

### Impact of VRFs on baseline RSFC

At baseline, we found no associations between VRFs and network or global functional connectivity (all p’s≥0.05), except that higher HDL-cholesterol levels were associated with lower baseline RSFC within the DMN (Schaefer, β=-0.146, p=0.023; Power, β=-0.141, p=0.028) and DAN network (Power, β=-0.129, p=0.045, see **eTable 1**).

### Global and network specific longitudinal change in RSFC across all individuals

We examined if RSFC changes over time without testing for an effect of VRFs. We found an overall increase in global RSFC over time when adjusted for age, sex and mean FD (Schaefer: β=0.009, s.e.=0.003, t=2.97, p=0.003; black dotted lines in **Figure 3**). Similar results were obtained when using the MIST and Power parcellation (MIST: β=0.009, s.e.=0.003, t=3.00, p=0.003; Power: β=0.010, s.e.=0.002, t=2.48, p=0.014; **eFigure 2**). No significant changes in DMN RSFC were observed over time across all individuals and all 3 parcellations (all p’s≥0.05). An overall increase in RSFC was found within specific functional networks across all individuals (Schaefer: in LIM and DAN, β’s ≥0.016, s.e.’s ≤0.006, t’s ≥2.97 p’s <0.003; MIST: in FPN and LIM, β’s ≥0.010, s.e.’s ≤0.004, t’s ≥2.09 p’s ≤0.037; Power: in DAN and SM, β’s ≥0.011, s.e.’s ≤0.007, t’s ≥2.05 p’s ≤0.040; **eFigure 2**).

**Figure 2:**
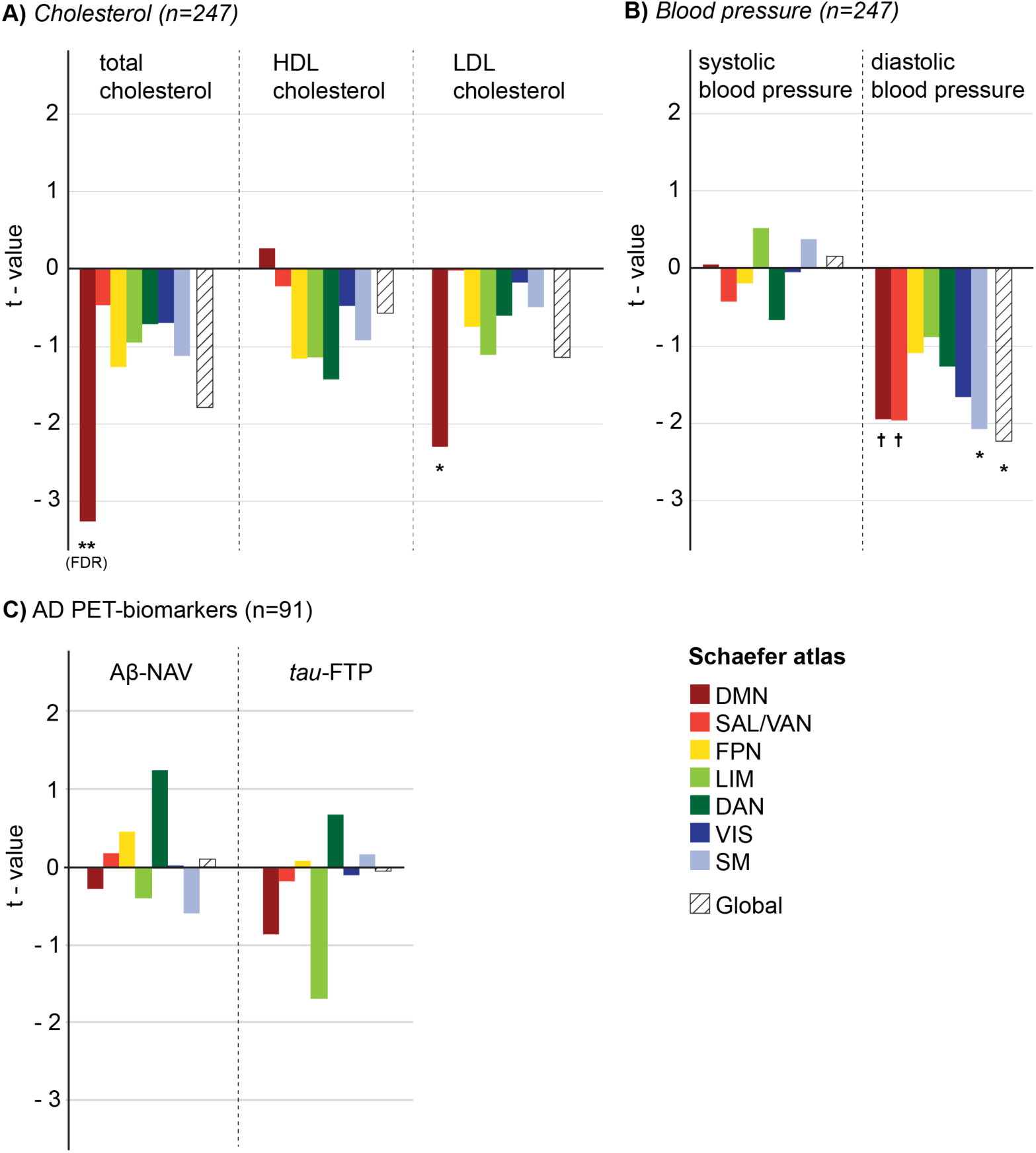
Summary of associations of VRF and AD biomarkers with functional connectivity changes over time from linear mixed effect models. T-values represent effects of (A) total-, HDL- and LDL-cholesterol, (B) systolic and diastolic blood pressure as well as (C) β-amyloid- (Aβ) and *tau*-PET SUVR on each RSFC network and on global RSFC, using the Schaefer parcellation atlas. Regarding vascular risk factors, mixed effects models were run either with total-cholesterol, HDL-cholesterol, systolic and diastolic blood pressure (Model A) or with LDL-cholesterol, HDL-cholesterol, systolic and diastolic blood pressure (Model B) as independent variables. For better visualisation, we averaged the t-values of both models for HDL-cholesterol and systolic and diastolic blood pressure, since they were highly similar (see eTable 2-4 for separate statistics). DAN, dorsal attention network; DMN, default mode network; FPN, fronto-parietal network; FTP, flortaucipir tracer; LIM, limbic network; NAV, Navidea tracer; SAL, salience network; SM, somatomotor network; VAN, ventral attention network and VIS, visual network. ^**†**^ p<0.05 only in model A or B, * p<0.05, ** p<0.01. Results that survived correction for false discovery rate are indicated by FDR.

**Figure 3:**
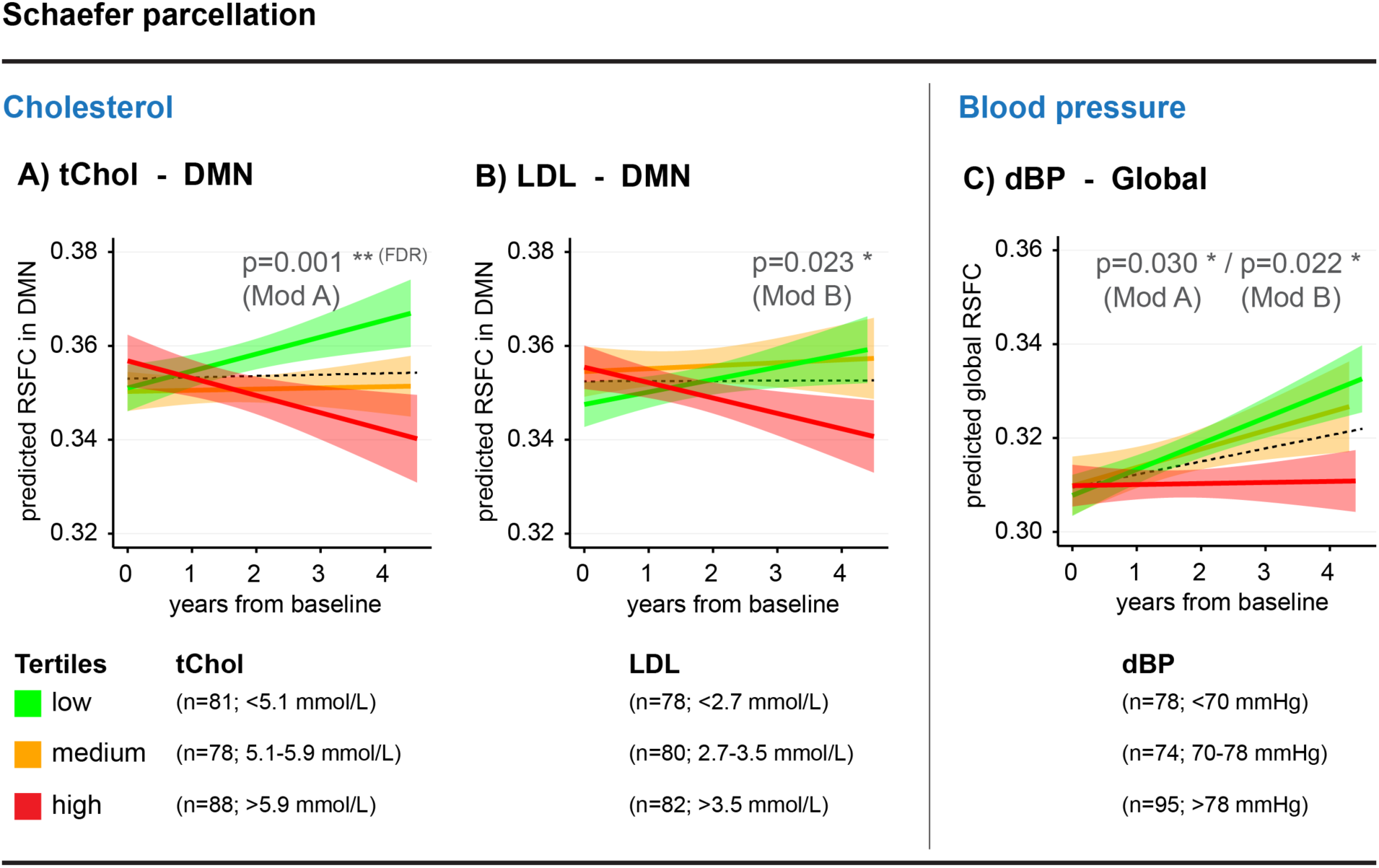
Change in resting-state functional connectivity, assessed using the Schaefer brain parcellation, as a function of vascular risk factors. These graphs are a longitudinal representation of a subset of the results shown in Figure 2. Results were only plotted when significant results were replicated with at least one other parcellation (MIST or Power). For visualisation purposes, we divided the continuous measures of (A) total-cholesterol, (B) LDL-cholesterol and (C) diastolic blood pressure into tertiles. Predicted RSFC estimates from linear-mixed effects models were plotted against follow-up time from baseline. Shaded regions represent 95% confidence intervals. Results that survived correction for false discovery rate are indicated by FDR. The dotted black lines represent the mean change in RSFC across all individuals. Total- and LDL-cholesterol exhibit a high multicollinearity, so separate models were run including one or the other (“total-cholesterol Model A” and the “LDL-cholesterol Model B”). All other vascular measures, and their interaction with time, were as well included in both models as predictors of RSFC changes. For better visualisation, graphics of diastolic blood pressure results were shown only for Model A, but statistics are presented from both Model A and B. dBP, diastolic blood pressure; DMN, default mode network; LDL, low-density lipoprotein; Mod, Model; RSFC, resting-state functional connectivity; tChol, total-cholesterol.

### Impact of VRFs on longitudinal change in RSFC

Using longitudinal analyses, we found no associations between any of the cholesterol values and global connectivity changes over time (Schaefer, all p’s≥0.05; **Figures 2** and **3, eTable 2**). Higher total-cholesterol levels were associated with decreased functional connectivity within the DMN (Schaefer, β=-0.013, s.e.=0.004, t=-3.26, p=0.001; **Figures 2** and **3, eTable 2**). This association survived FDR-correction and was similar when using the MIST and Power brain parcellations (**eFigures 1** and **2, eTables 3** and **4**). When the study sample was restricted to PET participants, this association was attenuated to trend-level (n=91, Schaefer, ß=-0.005, s.e.=0.003, t=-1.86, p=0.064). Higher total-cholesterol was also related to a reduction in RSFC within the SAL network; however, this was only seen with the Power parcellation atlas (**eFigures 1** and **2, eTable 4**). Higher LDL-cholesterol levels were associated with a decline in DMN RSFC (Schaefer, β=-0.010, s.e.=0.004, t=-2.30, p=0.023; **Figures 2** and **3, eTable 2**). This was also shown with the MIST parcellation, and at trend-level with the Power parcellation (**eFigures 1** and **2, eTables 3** and **4**). This association was attenuated when restricting analysis to PET participants (n=91, Schaefer, β=-0.004, s.e.=0.002, t=-1.45, p=0.138). No longitudinal effects on RSFC were found for HDL-cholesterol.

With regard to blood pressure, higher diastolic blood pressure levels were associated with a reduction of global RSFC over time (Mean Model A and B; Schaefer, β=-0.010, s.e.=0.004, t=-2.24, p=0.026; **Figures 2** and **3, eTable 2**). This association was similar when using the MIST and Power brain parcellations (**eFigures 1** and **2, eTables 3** and **4**). This association reached trend-level, when the study sample was restricted to the PET participants (n=91, Schaefer, ß=-0.004, s.e=0.002, t=-1.68, p=0.095). At the network level, associations were also found between higher diastolic blood pressure and reduction in RSFC within the DMN (Model A only; Schaefer, β=-0.008, s.e.=0.004, t=-2.00, p=0.047), SAL/VAN (Model A only; Schaefer, β=-0.010, s.e.=0.005, t=-1.98, p=0.049) and SM networks (Mean Model A and B; Schaefer, β=-0.017, s.e.=0.008, t=-2.07, p=0.040) (**Figure 2, eFigure 2** and **eTable 2**). The association between diastolic blood pressure and SAL/VAN RSFC was similar when using the MIST parcellations (**eFigures 1** and **2, eTable 3**). No associations were found between systolic blood pressure and change in RSFC. All detailed statistical results can be found in **eTables 2-4**.

### Impact of AD-related biomarkers on longitudinal change in RSFC

No association were found between global cerebral Aβ deposition or *tau* deposition in the entorhinal cortex and RSFC changes over time, using all three parcellations (p’s≥0.05; **eTable 5-7**).

## 4 Discussion

We investigated the impact of cholesterol and blood pressure on longitudinal RSFC changes within a well-established set of predefined functional networks covering the entire brain. The current study provides evidence that vascular burden contributes to RSFC trajectories in middle-to-late life adulthood. While RSFC was quite stable or slightly increasing across a 4-year follow-up (IQR 2-4 years), higher total- and LDL-cholesterol levels were associated with reductions in DMN RSFC over time, whereas higher diastolic blood pressure was associated with reduced whole-brain RSFC. In contrast, AD-related biomarkers showed no association with RSFC changes over time. These last analyses were unfortunately only performed in a subset of the cohort and will therefore need to be replicated. Given that decreased RSFC is usually associated with older age and clinical AD, VRFs may impair brain integrity independently and likely prior to AD pathology. In addition of being a risk factor for AD, some evidences suggest that vascular brain changes might precede AD-specific biomarker changes.^1^

We found an association of higher total- and LDL-cholesterol levels with a reduction in RSFC specifically within the DMN, whereas other functional networks showed no consistent changes in RSFC linked to cholesterol levels. The DMN has been suggested to have the highest metabolic rate in the brain, and to be involved in a large variety of demanding cognitive tasks.^27,46-48^ This high economical load may render the DMN preferentially vulnerable to aging and AD in comparison to other functional networks.^18,49^ Elevated vascular risk, measured via composite VRF measures, has been associated with a reduction in glucose metabolism,^50^ resting cerebral blood flow^51^ and cerebrovascular reactivity in DMN regions.^52^ Those vascular effects might evolve gradually in the presence of chronically elevated cholesterol levels and lead to disturbed brain functions. In comparison to other networks, RSFC seem to be related to cerebrovascular integrity particularly within the DMN.^53,54^ In line with this, a decrease in RSFC^54^ and functional connectivity density^55^ in the DMN has been found in patients with mild cognitive impairment with moderate to severe white matter hyperintensities, a measure of vascular brain integrity.

A reduction of DMN RSFC with advancing age has been recently shown in a longitudinal study with healthy older adults aged 60 years and older,^56^ a finding that was not replicated in our cohort. In contrast, our data suggests no change of DMN RSFC over time in the general cohort. When dividing the individuals by VRF profile, we found rather an increased or no change in DMN RSFC in individuals with lower to moderate cholesterol levels, whereas a decline in DMN RSFC was seen in the presence of high total- and LDL-cholesterol burden (see low, medium and high tertiles slopes in **Figure 3**). These differential trajectories highlight the relevant contribution of high VRFs to detect a decline in RSFC. Decline in DMN RSFC have been repetitively associated with clinical AD^10^, it is therefore possible that VRFs increase vulnerability to AD by creating early brain alteration in individuals at risk of AD.

Researchers have recently begun to investigate not only network-based RSFC, but also global RSFC (i.e. connectivity of each grey matter region with all other grey matter regions), as both seem to represent complementary predictors of AD pathologies and their propagation.^9,12^ Brain regions with high global RSFC are proposed to represent hub regions in the brain, mainly but not all of which are located within the DMN, showing especially high metabolic activity, neuronal activity and vulnerability to AD pathologies.^9,57^ While higher cholesterol levels showed network-specific associations with reduction in RSFC, higher diastolic, but not systolic, blood pressure was associated with reduction of global RSFC over time. Diastolic hypertension may specifically accelerate age-related sclerotic changes in small arterioles spread across the brain,^58^ leading to reduced cerebral perfusion, white matter lesions, atrophy and finally cognitive impairment.^59-61^ It has been also suggested that diastolic blood pressure is associated earlier on with structural impairment than systolic blood pressure; a similar time course may appear for functional impairment.^62^

Finally, we found increases in network and global RSFC over time across all individuals, without taking VRF differences into account. This hyperconnectivity has been reported in previous studies^9,20,63^ and has been typically interpreted as compensatory mechanisms to maintain cognitive function that starts to be compromised with aging and early AD.^64^ Newer concepts posit that neurons in the DMN might be unable to handle costly metabolic alterations due to higher pathology-induced processing burden, leading to a transfer of processing burden to downstream regions and/or noisy inefficient synaptic communications, which results in a short-term hyperconnectivity.^13,63^ Particularly, most central and metabolically active hub regions are likely to express increased connectivity.^63^ Increases in RSFC have been proposed to occurs until individuals reach an age of ∼74 years, after which a decline in RSFC begins.^9^ Our study with relatively young older adults (mean age of 63 years) suggests that such a cascading network failure might be accelerated in the presence of VRFs, by preluding the tipping point where high VRF burden overwhelms neurons of hub regions, resulting in functional disconnection.

In contrast to our consistent VRF findings and to previous AD-related studies^18,65^, we did not find associations of Aβ and *tau* deposition with changes in RSFC in our cohort at risk for AD. One can hypothesize that VRF may predate AD pathology, increase brain vulnerability and/or contribute to AD pathogenesis^22,66^; a hypothesis that needs further investigations. Our relatively small sample size might also have precluded us to identify mild associations. It will for instance be of high interest to replicate the association between *tau* and the limbic network (Schaefer, t-value = -1.785), the network that best overlap *tau* pathology, in a larger sample of older adults with more severe pathology.

A strength of our study is its longitudinal design to assess trajectories of functional connectivity in the context of vascular- and AD-related burden. Since RSFC was measured prospectively from the time point of VRF assessment, it can be hypothesised that VRFs may induce changes in RSFC; although this needs to be proven by future interventional trials. The reproduction of our results, using 3 different parcellation atlases, further strengthens their validity. There are also several caveats that must be taken into consideration when interpreting our findings. First, blood pressure values were assessed only from one measurement instead of averaging values of multiple measurements at baseline. Second, we used peripheral measures to assess cholesterol and blood pressure levels. 24S-hydroxycholesterol (the predominant metabolite of brain cholesterol), cerebral perfusion pressure or measures of cerebrovascular impairment (i.e. white matter lesions, lacunes, and microbleeds) may provide more direct evidence about a relationship between vascular burden, vascular brain injuries and changes in RSFC. Third, vascular burden was expressed only by elevated levels of cholesterol and blood pressure in our otherwise healthy cohort. The impact of other common VRFs, such as diabetes, smoking and obesity, still needs to be assessed in future studies. Nevertheless, the presence of VRFs was in general comparable to the US population,^67^ except that current smoking was on average 10-fold lower in the PREVENT-AD cohort, suggesting that our results are extendible to further populations. Fourth, it needs to be further explored if the observed VRF-related changes in RSFC reflect changes in blood flow, changes in neuronal activity or both.^68^ Fifth, levels of cerebral AD-related PET biomarkers were low to moderate in our relatively young cohort and the sample size was limited, therefore, associations of Aβ and *tau* with longitudinal RSFC may have been harder to detect.

In sum, our results suggest a significant contribution of VRFs to changes in DMN and global RSFC in cognitively normal older adults. This change seems to precede the effect of AD pathology in RSFC. In our cohort that includes only relatively young cognitively normal older adults with an elevated risk to develop AD, we found no effect of Aβ or *tau* on RSFC change suggesting this effect might only be capture later in the course of the disease. It will be of interest to follow these individuals over time to assess when AD pathology starts to impair RSFC and if VRFs and Aβ/*tau* burden have an additive or a synergistic effect on brain connectivity.

## Acknowledgments

This study was funded by the Alzheimer Society of Canada (*SV*), Brain Canada (*SV*), Quebec Bio-Imaging Network (*MM*), Healthy Brains for Healthy Lives (*MM*), the Alzheimer’s Association (*SV*), McGill University (*JB, JP*), the Fonds de Recherche du Québec – Santé (FRQ-S) (*JB, JP*), an unrestricted research grant from Pfizer Canada (*JB, JP*), the Levesque Foundation (*JP*), the Douglas Hospital Research Centre and Foundation (*JB, JP*), the Canada Institutes of Health Research (*SV*), the Canada Fund for Innovation (*SV*) and the German Research Foundation (DFG) (*TK*). The authors wish to acknowledge the PREVENT-AD staff, especially Jennifer Tremblay-Mercier, Joanne Frenette, Leslie-Ann Daoust, and the Brain Imaging Center of the Douglas Mental Health Research Institute, and the PET and cyclotron units of the Montreal Neurological Institute. A full listing of the PREVENT-AD Research Group members can be found at https://preventad.loris.ca/acknowledgements/acknowledgements.php?date=[2018-11-14]. We would also like to acknowledge the participants of the PREVENT-AD cohort for dedicating their time and energy to helping us collect these data.

## Author Contributions

**Theresa Köbe**: study concept and design, analysis and interpretation of data, drafting and revising the manuscript for intellectual content; **Alexa Pichet Binette**: data acquisition and analysis, revising the manuscript for intellectual content; **Jacob W. Vogel**: data analysis, revising the manuscript for intellectual content; **Pierre-François Meyer**: data analysis, revising the manuscript for intellectual content; **John Breitner**: acquisition of data, protocol concept and design, revising the manuscript for intellectual content; **Judes Poirier**: acquisition of data, protocol concept and design, revising the manuscript for intellectual content; **Sylvia Villeneuve**: acquisition of data, study concept and design, interpretation of data, revision of manuscript for intellectual content.

