## Supplementary Material for "Vascular burden is associated with a decline in default-mode and global resting-state functional connectivity in individuals at risk for Alzheimer’s disease"

---

#### eFigures

**eFigure 1:** Summary of associations of VRF and AD biomarkers with functional connectivity changes over time from linear mixed effect models

**eFigure 2:** Change in resting-state functional connectivity, assessed using 3 different brain parcellation atlases, as a function of vascular risk factors

#### eTables

**eTable 1:** Association between baseline VRF and RSFC at the first valid time point, using the Schaefer, MIST and Power parcellation

**eTable 2.** Association between VRF and longitudinal change in RSFC, using the Schaefer parcellation

**eTable 3.** Association between VRF and longitudinal change in RSFC, using the MIST parcellation

**eTable 4.** Association between VRF and longitudinal change in RSFC, using the Power parcellation

**eTable 5.** Association between AD biomarkers and longitudinal changes in RSFC, using the Schaefer parcellation

**eTable 6.** Association between AD biomarkers and longitudinal changes in RSFC, using the MIST parcellation

**eTable 7.** Association between AD biomarkers and longitudinal changes in RSFC, using the Power parcellation

### Supplementary methods

#### VRF assessment

All venous blood samples were taken non-fasting at enrollment, because most of each person's lifetime is spent in the postprandial state,[1] lipid profiles change minimally in response to normal food intake, and non-fasting lipid profiles seem to predict increased risk of cardiovascular events better than fasting lipid profiles.[2,3]
Note that new guidelines move towards a consensus on measuring lipid profiles for cardiovascular risk prediction in the non-fasting state.[3] In addition, only triglycerides and LDL-cholesterol (calculated based on triglyceride concentrations, using the Friedewald equation) seem to be influenced by non-fasting state. In the current study, however, triglycerides were not included, and LDL-cholesterol was measured using a direct homogeneous assay.

#### PET assessment

PET scans (amyloid-β, [$^{18}$F]NAV4694 (NAV) provided by Navidea Biopharmaceuticals (Dublin, Ohio) and *tau*, [$^{18}$F]AV1451 (Flortaucipir) provided by Eli Lilly & Company (Indianapolis, Indiana)) were acquired ordinarily on two consecutive days. Approximately 6mCi of NAV and 10mCi of Flortaucipir were injected intravenously. Static acquisition frames were obtained for Aβ at 40-70min (6x5min frames) and for *tau* at 80-100min (4x5min frames) post-injection.

#### APOE Genotyping

*APOE* genotype was determined using the PyroMark Q96 pyrosequencer (Qiagen, Toronto, ON, Canada) and the following primers: rs429358_amplification_forward 5'-ACGGCTGTCCAAGGAGCT G-3', rs429358_amplification_reverse_biotinylated 5'-CACCTCGCCGCGGTACTG-3', rs429358_sequencing 5'-CGGACATGGAGGACG-3', rs7412_amplification_forward 5'-CTCCGCGATGCCGATGAC-3', rs7412_amplification_reverse_biotinylated 5'-CCCCGGCCTGGTACACTG-3' and rs7412_sequencing 5'-CGATGACCTGCAGAAG-3'; also described previously.[4]

### Supplementary figures

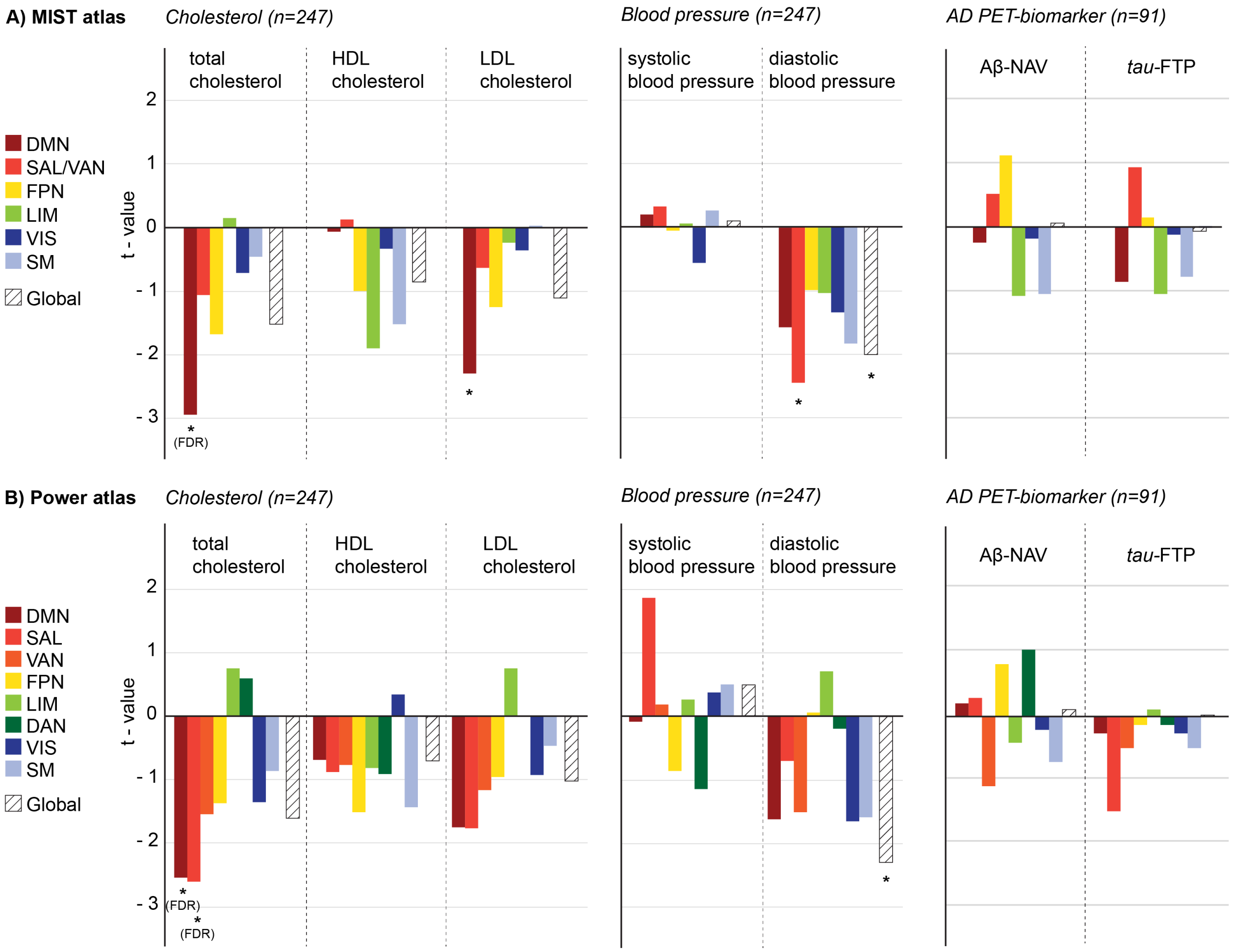


**eFigure 1: Summary of associations of VRF and AD biomarkers with functional connectivity changes over time from linear mixed effect models.** T-values represent effects of total-, HDL- and LDL-cholesterol, systolic and diastolic blood pressure as well as β-amyloid- (Aβ) and *tau*-PET SUVRs on each RSFC network and on global RSFC, using the MIST (A) and Power (B) parcellation atlases. Regarding vascular risk factors, linear-mixed effects models were run either with total-cholesterol, HDL-cholesterol, systolic and diastolic blood pressure as independent variables (Model A) or with LDL-cholesterol, HDL-cholesterol, systolic and diastolic blood pressure as independent variables (Model B). For better visualisation, we averaged the t-values of both models for HDL-cholesterol and systolic and diastolic blood pressure, since they were highly similar (see eTable 2-4 for separate statistics). DAN, dorsal attention network; DMN, default mode network; FPN, fronto-parietal network; FTP, flortaucipir tracer; LIM, limbic network; NAV, Navidea tracer; SAL, salience network; SM, somatomotor network; VAN, ventral attention network and VIS, visual network. * $p<0.05$. Results that survived correction for false discovery rate are indicated by FDR.

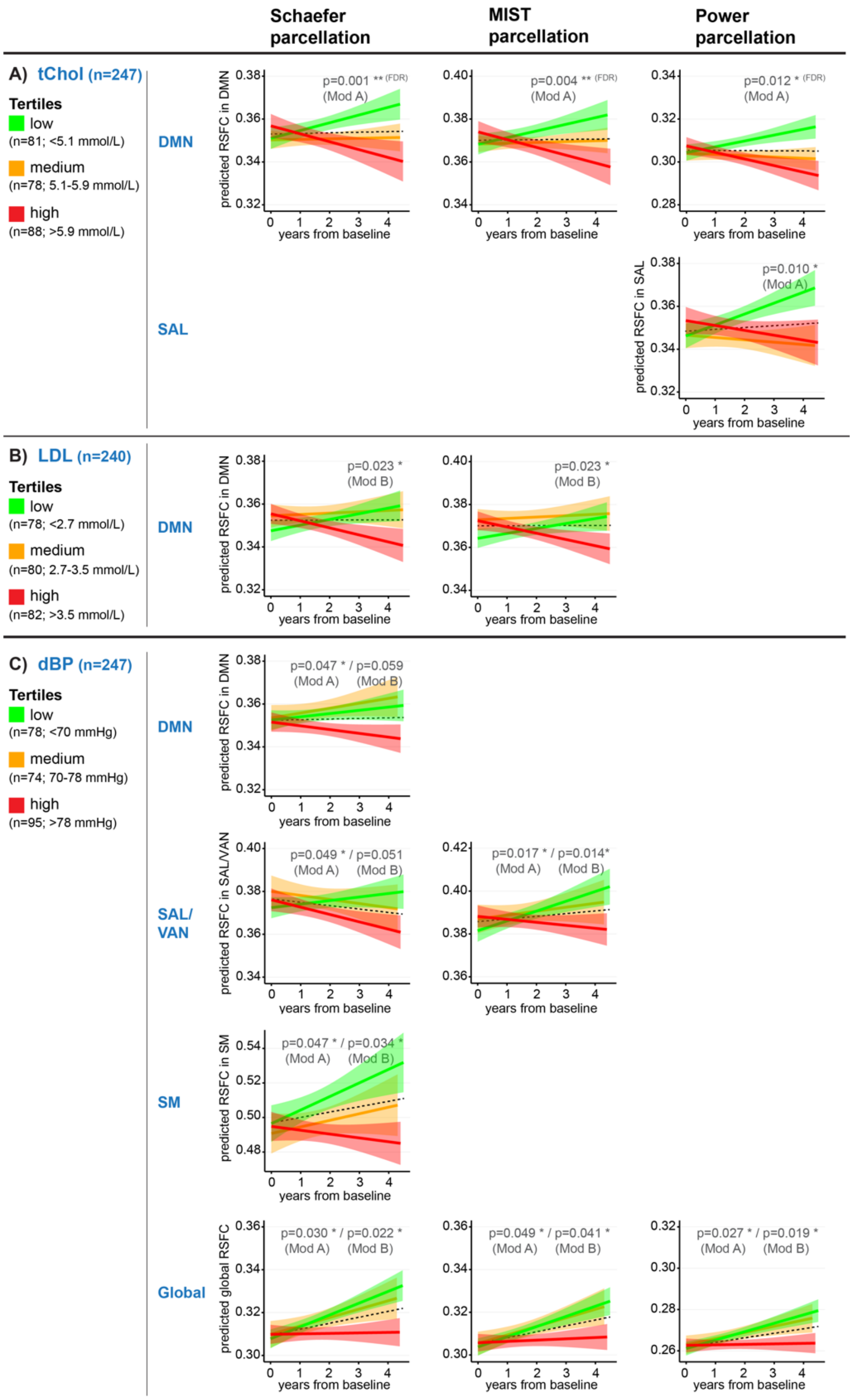


**eFigure 2: Change in resting-state functional connectivity, assessed using 3 different brain parcellation atlases, as a function of vascular risk factors.** These graphs represent a longitudinal representation of all significant results shown in Figure 2 of the main manuscript and eFigure 1. For visualisation purposes, we divided the continuous measures of total cholesterol (A), LDL cholesterol (B) and diastolic blood pressure (C) into tertiles. Predicted functional connectivity estimates from linear-mixed effects models were plotted against follow-up time from baseline. Shaded regions represent 95% confidence intervals and p-values are displayed as <0.01** and <0.05*. Results that survived correction for false discovery rate are indicated by FDR. The dotted black lines represent the mean change in RSFC of the entire sample (n=247). Diastolic blood pressure was included as predictor variable in both the "total-cholesterol Model A" and the "LDL-cholesterol Model B". Graphics in panel C were derived from Model A, but statistics are presented from both Models. dBP, diastolic blood pressure; DMN, default mode network; RSFC, resting-state functional connectivity; SAL/VAN, salience and ventral attention network; SM, somatomotor network; tChol, total cholesterol.

### Supplementary tables

##### eTable 1. Association between baseline vascular risk factors and RSFC at the first valid time point, using the Schaefer, MIST and Power parcellation

| Schaefer parcellation | | DMN | SAL/VAN | | FPN | LIM | DAN | VIS | SM | global |
|---|---|---|---|---|---|---|---|---|---|---|
| **tChol** | estimate | 0.060 | 0.111 | | 0.081 | 0.030 | 0.064 | 0.026 | 0.053 | 0.100 |
| n=247 | p-value | 0.348 | 0.085 | | 0.210 | 0.644 | 0.322 | 0.688 | 0.412 | 0.118 |
| **HDL** | estimate | -0.146 | -0.072 | | -0.097 | -0.036 | -0.075 | -0.049 | -0.049 | -0.102 |
| n=247 | p-value | **0.023*** | 0.267 | | 0.132 | 0.578 | 0.244 | 0.446 | 0.443 | 0.115 |
| **LDL** | estimate | 0.093 | 0.113 | | 0.084 | 0.019 | 0.085 | 0.018 | 0.045 | 0.115 |
| n=240 | p-value | 0.155 | 0.123 | | 0.199 | 0.769 | 0.196 | 0.781 | 0.487 | 0.077 |
| **sBP** | estimate | -0.053 | 0.009 | | -0.019 | -0.080 | -0.038 | -0.018 | -0.054 | -0.039 |
| n=247 | p-value | 0.414 | 0.893 | | 0.773 | 0.211 | 0.560 | 0.783 | 0.398 | 0.543 |
| **dBP** | estimate | 0.012 | 0.066 | | 0.042 | 0.014 | 0.037 | -0.049 | -0.008 | 0.047 |
| n=247 | p-value | 0.854 | 0.303 | | 0.517 | 0.827 | 0.563 | 0.447 | 0.905 | 0.467 |

| MIST parcellation | | DMN | SAL/VAN | | FPN | LIM | | VIS | SM | global |
|---|---|---|---|---|---|---|---|---|---|---|
| **tChol** | estimate | 0.051 | 0.066 | | 0.088 | 0.068 | | 0.024 | 0.033 | 0.097 |
| n=247 | p-value | 0.429 | 0.304 | | 0.174 | 0.292 | | 0.708 | 0.609 | 0.132 |
| **HDL** | estimate | -0.112 | -0.122 | | -0.108 | 0.085 | | -0.061 | -0.033 | -0.084 |
| n=247 | p-value | 0.082 | 0.058 | | 0.094 | 0.189 | | 0.347 | 0.610 | 0.194 |
| **LDL** | estimate | 0.086 | 0.102 | | 0.115 | 0.074 | | 0.019 | 0.008 | 0.118 |
| n=240 | p-value | 0.189 | 0.119 | | 0.078 | 0.257 | | 0.773 | 0.906 | 0.070 |
| **sBP** | estimate | -0.082 | 0.017 | | -0.031 | -0.075 | | 0.043 | -0.038 | -0.028 |
| n=247 | p-value | 0.202 | 0.788 | | 0.634 | 0.244 | | 0.501 | 0.554 | 0.665 |
| **dBP** | estimate | 0.022 | 0.103 | | 0.040 | 0.022 | | -0.039 | -0.025 | 0.055 |
| n=247 | p-value | 0.732 | 0.109 | | 0.539 | 0.737 | | 0.541 | 0.694 | 0.390 |

| Power parcellation | | DMN | SAL | VAN | FPN | LIM | DAN | VIS | SM | global |
|---|---|---|---|---|---|---|---|---|---|---|
| **tChol** | estimate | 0.028 | 0.088 | 0.032 | 0.061 | 0.026 | 0.036 | 0.025 | 0.020 | 0.071 |
| n=247 | p-value | 0.661 | 0.173 | 0.614 | 0.346 | 0.685 | 0.572 | 0.693 | 0.752 | 0.271 |
| **HDL** | estimate | -0.141 | 0.005 | -0.110 | -0.046 | 0.120 | -0.129 | -0.045 | -0.019 | -0.083 |
| n=247 | p-value | **0.028*** | 0.943 | 0.087 | 0.476 | 0.061 | **0.045*** | 0.488 | 0.766 | 0.200 |
| **LDL** | estimate | 0.064 | 0.100 | 0.085 | 0.085 | 0.005 | 0.116 | 0.018 | -0.016 | 0.100 |
| n=240 | p-value | 0.329 | 0.125 | 0.193 | 0.192 | 0.934 | 0.075 | 0.782 | 0.804 | 0.127 |
| **sBP** | estimate | -0.029 | -0.089 | -0.055 | 0.055 | -0.034 | -0.067 | 0.031 | -0.057 | -0.031 |
| n=247 | p-value | 0.651 | 0.165 | 0.397 | 0.390 | 0.602 | 0.295 | 0.626 | 0.378 | 0.631 |
| **dBP** | estimate | 0.014 | -0.042 | 0.010 | 0.096 | -0.057 | -0.071 | -0.045 | -0.020 | 0.030 |
| n=247 | p-value | 0.829 | 0.510 | 0.880 | 0.134 | 0.375 | 0.269 | 0.490 | 0.757 | 0.647 |

General linear models were performed to test for associations of total-cholesterol, HDL-cholesterol, LDL-cholesterol, systolic blood pressure and diastolic blood pressure with resting-state functional connectivity (RSFC) at the first time point with a valid resting-state scan (n=240 at baseline, n=5 at follow-up year 1, n=2 at follow-up year 2). RSFC was estimated within the default mode (DMN), salience and ventral attention (SAL/VAN), fronto-parietal (FPN), limbic (LIM), dorsal attention (DAN), visual (VIS) and somatomotor network (SM). Correlation analyses were corrected for age, sex, use of vascular medication and mean framewise displacement. * $p<0.05$.

#### eTable 2. Association between VRF and longitudinal change in RSFC, using the Schaefer parcellation

| Model A | DMN | | | | SAL/ VAN | | | | FPN | | | | LIM | | | |
|---|---|---|---|---|---|---|---|---|---|---|---|---|---|---|---|---|
| n of obs. = 865<br>n of part. = 247 | Est. | SE | t | p | Est. | SE | t | p | Est. | SE | t | p | Est. | SE | t | p |
| tChol x time | -0.0126 | 0.004 | -3.263 | 0.001** | -0.0023 | 0.005 | -0.464 | 0.643 | -0.0056 | 0.004 | -1.270 | 0.204 | -0.0073 | 0.008 | -0.947 | 0.345 |
| HDL x time | 0.0032 | 0.004 | 0.761 | 0.447 | -0.0011 | 0.005 | -0.207 | 0.836 | -0.0042 | 0.005 | -0.893 | 0.372 | -0.0081 | 0.008 | -0.986 | 0.325 |
| sBP x time | 0.0008 | 0.004 | 0.182 | 0.856 | -0.0026 | 0.006 | -0.459 | 0.647 | -0.0009 | 0.005 | -0.175 | 0.861 | 0.0050 | 0.009 | 0.567 | 0.572 |
| dBP x time | -0.0080 | 0.004 | -1.997 | 0.047* | -0.0099 | 0.005 | -1.976 | 0.049* | -0.0055 | 0.005 | -1.217 | 0.224 | -0.0069 | 0.008 | -0.867 | 0.387 |
| age x time | -0.0024 | 0.003 | -0.773 | 0.441 | 0.0078 | 0.004 | 1.959 | 0.052. | -0.0032 | 0.004 | -0.899 | 0.369 | 0.0045 | 0.006 | 0.710 | 0.479 |
| sex x time | -0.0018 | 0.008 | -0.225 | 0.822 | -0.0047 | 0.010 | -0.475 | 0.635 | 0.0059 | 0.009 | 0.662 | 0.508 | 0.0385 | 0.016 | 2.467 | 0.015* |
| medication x time | -0.0020 | 0.007 | -0.273 | 0.785 | 0.0001 | 0.009 | 0.009 | 0.993 | 0.0067 | 0.008 | 0.788 | 0.431 | -0.0037 | 0.015 | -0.251 | 0.802 |
| mean FD x time | 0.0112 | 0.003 | 3.214 | 0.001** | 0.0074 | 0.004 | 1.728 | 0.085. | 0.0054 | 0.004 | 1.370 | 0.171 | 0.0087 | 0.007 | 1.283 | 0.201 |
| tChol | 0.0017 | 0.003 | 0.628 | 0.531 | 0.0072 | 0.003 | 2.326 | 0.021* | 0.0052 | 0.003 | 1.517 | 0.131 | 0.0017 | 0.007 | 0.261 | 0.794 |
| HDL | -0.0079 | 0.003 | -2.894 | 0.004** | -0.0072 | 0.003 | -2.349 | 0.020* | -0.0096 | 0.003 | -2.803 | 0.005** | -0.0081 | 0.007 | -1.234 | 0.218 |
| sBP | -0.0011 | 0.003 | -0.355 | 0.723 | -0.0023 | 0.003 | -0.675 | 0.500 | -0.0036 | 0.004 | -0.953 | 0.342 | -0.0077 | 0.007 | -1.060 | 0.290 |
| dBP | -0.0033 | 0.003 | -1.181 | 0.239 | -0.0024 | 0.003 | -0.767 | 0.444 | 0.0003 | 0.003 | 0.078 | 0.938 | -0.0014 | 0.007 | -0.212 | 0.833 |
| age | 0.0026 | 0.002 | 1.131 | 0.259 | 0.0053 | 0.003 | 2.063 | 0.040* | -0.0009 | 0.003 | -0.296 | 0.768 | 0.0036 | 0.006 | 0.657 | 0.512 |
| sex | 0.0001 | 0.006 | 0.015 | 0.988 | -0.0162 | 0.006 | -2.565 | 0.011* | -0.0108 | 0.007 | -1.534 | 0.126 | -0.0172 | 0.014 | -1.264 | 0.207 |
| medication | -0.0079 | 0.005 | -1.532 | 0.127 | 0.0057 | 0.006 | 0.977 | 0.329 | 0.0043 | 0.007 | 0.649 | 0.517 | -0.0025 | 0.013 | -0.199 | 0.843 |
| mean FD | -0.0003 | 0.002 | -0.154 | 0.878 | -0.0038 | 0.002 | -1.807 | 0.071. | 0.0048 | 0.002 | 2.246 | 0.025* | 0.0211 | 0.004 | 5.703 | 0.000*** |
| time | 0.0013 | 0.007 | 0.178 | 0.859 | -0.0070 | 0.009 | -0.776 | 0.439 | -0.0020 | 0.008 | -0.247 | 0.805 | -0.0007 | 0.014 | -0.051 | 0.960 |
| R2m | 0.055 | | | | 0.071 | | | | 0.058 | | | | 0.084 | | | |
| R2c | 0.515 | | | | 0.518 | | | | 0.567 | | | | 0.689 | | | |

| Model A | DAN | | | | VIS | | | | SM | | | | global | | | |
|---|---|---|---|---|---|---|---|---|---|---|---|---|---|---|---|---|
| n of obs. = 865<br>n of part. = 247 | Est. | SE | t | p | Est. | SE | t | p | Est. | SE | t | p | Est. | SE | t | p |
| tChol x time | -0.0046 | 0.006 | -0.709 | 0.479 | -0.0053 | 0.008 | -0.697 | 0.486 | -0.0086 | 0.008 | -1.123 | 0.263 | -0.0066 | 0.004 | -1.793 | 0.073. |
| HDL x time | -0.0084 | 0.007 | -1.206 | 0.228 | -0.0031 | 0.008 | -0.388 | 0.698 | -0.0064 | 0.008 | -0.779 | 0.437 | -0.0012 | 0.004 | -0.315 | 0.753 |
| sBP x time | -0.0050 | 0.008 | -0.661 | 0.509 | -0.0010 | 0.009 | -0.117 | 0.907 | 0.0032 | 0.009 | 0.358 | 0.721 | 0.0004 | 0.004 | 0.095 | 0.924 |
| dBP x time | -0.0091 | 0.007 | -1.358 | 0.175 | -0.0117 | 0.008 | -1.503 | 0.133 | -0.0158 | 0.008 | -1.999 | 0.047* | -0.0083 | 0.004 | -2.175 | 0.030* |
| age x time | -0.0057 | 0.005 | -1.077 | 0.282 | -0.0015 | 0.006 | -0.246 | 0.806 | 0.0040 | 0.006 | 0.638 | 0.525 | -0.0006 | 0.003 | -0.189 | 0.850 |
| sex x time | 0.0172 | 0.013 | 1.308 | 0.191 | -0.0001 | 0.015 | -0.008 | 0.993 | 0.0289 | 0.016 | 1.853 | 0.066. | 0.0068 | 0.007 | 0.913 | 0.361 |
| medication x time | -0.0103 | 0.013 | -0.822 | 0.411 | -0.0097 | 0.015 | -0.669 | 0.504 | -0.0024 | 0.015 | -0.164 | 0.870 | -0.0023 | 0.007 | -0.328 | 0.743 |
| mean FD x time | 0.0020 | 0.006 | 0.338 | 0.735 | 0.0059 | 0.007 | 0.860 | 0.390 | 0.0036 | 0.007 | 0.529 | 0.597 | 0.0051 | 0.003 | 1.518 | 0.130 |
| tChol | 0.0075 | 0.005 | 1.606 | 0.110 | 0.0039 | 0.005 | 0.709 | 0.479 | 0.0065 | 0.005 | 1.197 | 0.233 | 0.0051 | 0.003 | 1.933 | 0.054. |
| HDL | -0.0110 | 0.005 | -2.355 | 0.019* | -0.0112 | 0.005 | -2.051 | 0.041* | -0.0113 | 0.005 | -2.083 | 0.038* | -0.0081 | 0.003 | -3.037 | 0.003** |
| sBP | -0.0067 | 0.005 | -1.306 | 0.193 | 0.0050 | 0.006 | 0.834 | 0.405 | 0.0001 | 0.006 | 0.010 | 0.992 | -0.0016 | 0.003 | -0.563 | 0.574 |
| dBP | -0.0010 | 0.005 | -0.211 | 0.833 | -0.0156 | 0.006 | -2.807 | 0.005** | -0.0087 | 0.006 | -1.589 | 0.114 | -0.0034 | 0.003 | -1.250 | 0.213 |
| age | 0.0015 | 0.004 | 0.382 | 0.703 | -0.0032 | 0.005 | -0.705 | 0.481 | 0.0011 | 0.005 | 0.244 | 0.808 | 0.0035 | 0.002 | 1.568 | 0.118 |
| sex | -0.0464 | 0.010 | -4.826 | 0.000*** | -0.0171 | 0.011 | -1.520 | 0.130 | -0.0447 | 0.011 | -4.008 | 0.000*** | -0.0141 | 0.005 | -2.587 | 0.010* |
| medication | 0.0082 | 0.009 | 0.918 | 0.360 | -0.0004 | 0.010 | -0.039 | 0.969 | 0.0043 | 0.010 | 0.415 | 0.679 | 0.0041 | 0.005 | 0.803 | 0.423 |
| mean FD | 0.0044 | 0.003 | 1.406 | 0.160 | 0.0022 | 0.004 | 0.610 | 0.542 | -0.0065 | 0.004 | -1.824 | 0.069. | 0.0015 | 0.002 | 0.875 | 0.382 |
| time | 0.0069 | 0.012 | 0.574 | 0.566 | 0.0040 | 0.014 | 0.290 | 0.772 | -0.0116 | 0.014 | -0.823 | 0.411 | 0.0039 | 0.007 | 0.569 | 0.569 |
| R2m | 0.114 | | | | 0.045 | | | | 0.087 | | | | 0.085 | | | |
| R2c | 0.543 | | | | 0.509 | | | | 0.566 | | | | 0.527 | | | |

| Model B | DMN | | | | SAL/ VAN | | | | FPN | | | | LIM | | | |
|---|---|---|---|---|---|---|---|---|---|---|---|---|---|---|---|---|
| n of obs = 838<br>n of part. = 240 | Est. | SE | t | p | Est. | SE | t | p | Est. | SE | t | p | Est. | SE | t | p |
| LDL x time | -0.0092 | 0.004 | -2.301 | 0.023* | -0.0023 | 0.005 | -0.464 | 0.643 | -0.0033 | 0.004 | -0.742 | 0.459 | -0.0087 | 0.008 | -1.106 | 0.270 |
| HDL x time | -0.0009 | 0.004 | -0.213 | 0.831 | -0.0011 | 0.005 | -0.207 | 0.836 | -0.0063 | 0.004 | -1.420 | 0.158 | -0.0102 | 0.008 | -1.290 | 0.199 |
| sBP x time | -0.0006 | 0.005 | -0.122 | 0.903 | -0.0026 | 0.006 | -0.459 | 0.647 | -0.0010 | 0.005 | -0.197 | 0.844 | 0.0043 | 0.009 | 0.474 | 0.636 |
| dBP x time | -0.0078 | 0.004 | -1.900 | 0.059. | -0.0099 | 0.005 | -1.976 | 0.050. | -0.0044 | 0.005 | -0.968 | 0.335 | -0.0074 | 0.008 | -0.907 | 0.366 |
| age x time | -0.0028 | 0.003 | -0.848 | 0.398 | 0.0078 | 0.004 | 1.959 | 0.052. | -0.0033 | 0.004 | -0.917 | 0.361 | 0.0037 | 0.006 | 0.575 | 0.567 |
| sex x time | -0.0043 | 0.008 | -0.520 | 0.604 | -0.0047 | 0.010 | -0.475 | 0.635 | 0.0049 | 0.009 | 0.544 | 0.587 | 0.0347 | 0.016 | 2.132 | 0.035* |
| medication x time | 0.0009 | 0.008 | 0.115 | 0.908 | 0.0001 | 0.009 | 0.009 | 0.993 | 0.0063 | 0.009 | 0.713 | 0.477 | -0.0046 | 0.016 | -0.289 | 0.773 |
| mean FD x time | 0.0097 | 0.004 | 2.754 | 0.006** | 0.0074 | 0.004 | 1.728 | 0.085 | 0.0043 | 0.004 | 1.094 | 0.275 | 0.0087 | 0.007 | 1.254 | 0.211 |
| LDL | 0.0025 | 0.003 | 0.955 | 0.341 | 0.0072 | 0.003 | 2.326 | 0.021** | 0.0041 | 0.003 | 1.237 | 0.217 | -0.0010 | 0.007 | -0.155 | 0.877 |
| HDL | -0.0072 | 0.003 | -2.793 | 0.006** | -0.0072 | 0.003 | -2.349 | 0.020. | -0.0078 | 0.003 | -2.374 | 0.018* | -0.0072 | 0.006 | -1.135 | 0.258 |
| sBP | 0.0000 | 0.003 | 0.007 | 0.994 | -0.0023 | 0.003 | -0.675 | 0.500 | -0.0024 | 0.004 | -0.625 | 0.533 | -0.0069 | 0.007 | -0.929 | 0.354 |
| dBP | -0.0041 | 0.003 | -1.492 | 0.137 | -0.0024 | 0.003 | -0.767 | 0.444 | 0.0003 | 0.004 | 0.075 | 0.940 | -0.0007 | 0.007 | -0.098 | 0.922 |
| age | 0.0027 | 0.002 | 1.215 | 0.226 | 0.0053 | 0.003 | 2.063 | 0.040* | -0.0009 | 0.003 | -0.317 | 0.752 | 0.0029 | 0.006 | 0.510 | 0.611 |
| sex | -0.0005 | 0.006 | -0.098 | 0.922 | -0.0162 | 0.006 | -2.565 | 0.011** | -0.0097 | 0.007 | -1.347 | 0.179 | -0.0163 | 0.014 | -1.166 | 0.245 |
| medication | -0.0062 | 0.005 | -1.176 | 0.241 | 0.0057 | 0.006 | 0.977 | 0.329 | 0.0036 | 0.007 | 0.528 | 0.598 | -0.0076 | 0.013 | -0.577 | 0.565 |
| mean FD | -0.0010 | 0.002 | -0.537 | 0.591 | -0.0038 | 0.002 | -1.807 | 0.071. | 0.0041 | 0.002 | 1.900 | 0.058. | 0.0218 | 0.004 | 5.812 | 0.000*** |
| time | 0.0024 | 0.008 | 0.315 | 0.753 | -0.0070 | 0.009 | -0.776 | 0.439 | -0.0008 | 0.008 | -0.095 | 0.925 | 0.0026 | 0.015 | 0.174 | 0.862 |
| R2m | 0.052 | | | | 0.078 | | | | 0.043 | | | | 0.081 | | | |
| R2c | 0.534 | | | | 0.526 | | | | 0.569 | | | | 0.698 | | | |

| Model B | DAN | | | | VIS | | | | SM | | | | global | | | |
|---|---|---|---|---|---|---|---|---|---|---|---|---|---|---|---|---|
| n of obs = 838<br>n of part. = 240 | Est. | SE | t | p | Est. | SE | t | p | Est. | SE | t | p | Est. | SE | t | p |
| LDL x time | -0.0039 | 0.007 | -0.596 | 0.552 | -0.0013 | 0.008 | -0.177 | 0.860 | -0.0039 | 0.008 | -0.495 | 0.621 | -0.0042 | 0.004 | -1.144 | 0.253 |
| HDL x time | -0.0109 | 0.007 | -1.637 | 0.104 | -0.0045 | 0.008 | -0.577 | 0.564 | -0.0084 | 0.008 | -1.062 | 0.289 | -0.0031 | 0.004 | -0.827 | 0.409 |
| sBP x time | -0.0052 | 0.008 | -0.677 | 0.499 | 0.0000 | 0.009 | 0.003 | 0.997 | 0.0037 | 0.009 | 0.400 | 0.690 | 0.0009 | 0.004 | 0.211 | 0.833 |
| dBP x time | -0.0080 | 0.007 | -1.188 | 0.237 | -0.0143 | 0.008 | -1.812 | 0.070. | -0.0174 | 0.008 | -2.138 | 0.034* | -0.0087 | 0.004 | -2.294 | 0.022* |
| age x time | -0.0065 | 0.005 | -1.221 | 0.224 | -0.0008 | 0.006 | -0.136 | 0.892 | 0.0050 | 0.006 | 0.777 | 0.438 | -0.0002 | 0.003 | -0.063 | 0.950 |
| sex x time | 0.0149 | 0.013 | 1.104 | 0.272 | -0.0044 | 0.016 | -0.282 | 0.778 | 0.0284 | 0.016 | 1.745 | 0.083. | 0.0056 | 0.008 | 0.741 | 0.459 |
| medication x time | -0.0134 | 0.013 | -1.014 | 0.312 | -0.0026 | 0.015 | -0.170 | 0.865 | 0.0032 | 0.016 | 0.202 | 0.840 | 0.0000 | 0.007 | -0.002 | 0.999 |
| mean FD x time | 0.0028 | 0.006 | 0.470 | 0.639 | 0.0051 | 0.007 | 0.736 | 0.462 | 0.0022 | 0.007 | 0.322 | 0.748 | 0.0036 | 0.003 | 1.099 | 0.272 |
| LDL | 0.0071 | 0.005 | 1.543 | 0.124 | 0.0049 | 0.005 | 0.917 | 0.360 | 0.0058 | 0.005 | 1.076 | 0.283 | 0.0050 | 0.003 | 1.926 | 0.055. |
| HDL | -0.0104 | 0.005 | -2.290 | 0.023* | -0.0097 | 0.005 | -1.849 | 0.066. | -0.0099 | 0.005 | -1.870 | 0.063. | -0.0066 | 0.003 | -2.625 | 0.009** |
| sBP | -0.0052 | 0.005 | -0.979 | 0.329 | 0.0060 | 0.006 | 0.984 | 0.326 | 0.0007 | 0.006 | 0.118 | 0.906 | -0.0005 | 0.003 | -0.186 | 0.853 |
| dBP | -0.0016 | 0.005 | -0.326 | 0.745 | -0.0162 | 0.006 | -2.887 | 0.004** | -0.0090 | 0.006 | -1.599 | 0.111 | -0.0039 | 0.003 | -1.451 | 0.148 |
| age | 0.0025 | 0.004 | 0.621 | 0.535 | -0.0026 | 0.005 | -0.560 | 0.576 | 0.0018 | 0.005 | 0.387 | 0.699 | 0.0039 | 0.002 | 1.730 | 0.085. |
| sex | -0.0471 | 0.010 | -4.761 | 0.000*** | -0.0149 | 0.011 | -1.299 | 0.195 | -0.0447 | 0.012 | -3.878 | 0.000*** | -0.0135 | 0.006 | -2.451 | 0.015* |
| medication | 0.0084 | 0.009 | 0.896 | 0.371 | 0.0006 | 0.011 | 0.054 | 0.957 | 0.0055 | 0.011 | 0.510 | 0.611 | 0.0050 | 0.005 | 0.960 | 0.338 |
| mean FD | 0.0042 | 0.003 | 1.351 | 0.177 | 0.0024 | 0.004 | 0.650 | 0.516 | -0.0062 | 0.004 | -1.725 | 0.085. | 0.0012 | 0.002 | 0.672 | 0.502 |
| time | 0.0107 | 0.012 | 0.862 | 0.390 | 0.0046 | 0.014 | 0.322 | 0.748 | -0.0143 | 0.015 | -0.960 | 0.338 | 0.0038 | 0.007 | 0.550 | 0.582 |
| R2m | 0.111 | | | | 0.041 | | | | 0.082 | | | | 0.077 | | | |
| R2c | 0.552 | | | | 0.508 | | | | 0.569 | | | | 0.528 | | | |

Using linear mixed effects models, we test whether vascular risk factors (VRF) were associated with a change in resting-state functional connectivity (RSFC) within different functional networks that were defined by the Schaefer parcellation atlas (DMN, default mode network; SAL/VAN, salience and ventral attention network; FPN, fronto-parietal network; LIM, limbic network; DAN, dorsal attention network; VIS, visual network and SM, somatomotor network) and throughout the whole brain (global). Two separate models were conducted, where Model A included total cholesterol (tChol), high-density lipoprotein cholesterol (HDL), systolic blood pressure (sBP) and diastolic blood pressure (dBP) and Model B included low-density lipoprotein cholesterol (LDL), HDL, sBP and dBP, due to the severe multicollinearity between tChol and LDL. As random effects intercept and slope of RSFC for each individual were included in the models. Models were corrected for baseline age, sex and vascular medication (intake or non-intake of drugs against dyslipidemia and/or hypertension), for longitudinal mean frame-displacement (FD) and for the interactions of those covariates with time. Unstandardized estimates (Est.), standard errors (SE), t-values and p-values (***p<0.001, **p<0.01, *p<0.05) as well as marginal (R2m) and conditional (R2c) $R^2$ values are presented. Obs., observations; part, participants.

#### eTable 3. Association between VRF and longitudinal change in RSFC, using the MIST parcellation

| Model A | DMN | | | | SAL/ VAN | | | | FPN | | | | LIM | | | | VIS | | | | SM | | | | global | | | |
|---|---|---|---|---|---|---|---|---|---|---|---|---|---|---|---|---|---|---|---|---|---|---|---|---|---|---|---|---|
| n of obs. = 865 | | | | | | | | | | | | | | | | | | | | | | | | | | | | |
| n of part. = 247 | Est. | SE | t | p | Est. | SE | t | p | Est. | SE | t | p | Est. | SE | t | p | Est. | SE | t | p | Est. | SE | t | p | Est. | SE | t | p |
| tChol x time | -0.0116 | 0.004 | -2.944 | 0.004** | -0.0048 | 0.005 | -1.053 | 0.294 | -0.0063 | 0.004 | -1.672 | 0.095. | 0.0008 | 0.005 | 0.157 | 0.876 | -0.0075 | 0.010 | -0.717 | 0.474 | -0.0057 | 0.012 | -0.459 | 0.647 | -0.0052 | 0.003 | -1.518 | 0.129 |
| HDL x time | 0.0018 | 0.004 | 0.423 | 0.673 | 0.0011 | 0.005 | 0.219 | 0.827 | -0.0028 | 0.004 | -0.697 | 0.486 | -0.0101 | 0.005 | -1.881 | 0.061. | -0.0032 | 0.011 | -0.287 | 0.775 | -0.0194 | 0.013 | -1.466 | 0.144 | -0.0023 | 0.004 | -0.628 | 0.530 |
| sBP x time | 0.0014 | 0.005 | 0.306 | 0.760 | 0.0016 | 0.005 | 0.312 | 0.755 | -0.0003 | 0.004 | -0.074 | 0.941 | 0.0003 | 0.006 | 0.060 | 0.953 | -0.0074 | 0.012 | -0.612 | 0.542 | 0.0046 | 0.014 | 0.323 | 0.747 | 0.0000 | 0.004 | 0.010 | 0.992 |
| dBP x time | -0.0065 | 0.004 | -1.618 | 0.107 | -0.0113 | 0.005 | -2.419 | 0.017* | -0.0040 | 0.004 | -1.046 | 0.296 | -0.0056 | 0.005 | -1.062 | 0.290 | -0.0129 | 0.011 | -1.199 | 0.232 | -0.0236 | 0.013 | -1.840 | 0.068. | -0.0069 | 0.004 | -1.972 | 0.049* |
| age x time | -0.0018 | 0.003 | -0.567 | 0.571 | 0.0034 | 0.004 | 0.908 | 0.365 | -0.0027 | 0.003 | -0.894 | 0.372 | 0.0005 | 0.004 | 0.113 | 0.910 | -0.0067 | 0.009 | -0.781 | 0.436 | 0.0001 | 0.010 | 0.006 | 0.996 | -0.0006 | 0.003 | -0.209 | 0.834 |
| sex x time | 0.0025 | 0.008 | 0.313 | 0.755 | 0.0097 | 0.009 | 1.051 | 0.295 | 0.0077 | 0.008 | 1.012 | 0.312 | 0.0133 | 0.010 | 1.286 | 0.200 | 0.0003 | 0.021 | 0.012 | 0.990 | 0.0360 | 0.025 | 1.427 | 0.156 | 0.0062 | 0.007 | 0.904 | 0.366 |
| medication x time | -0.0066 | 0.008 | -0.873 | 0.383 | -0.0025 | 0.009 | -0.290 | 0.772 | 0.0004 | 0.007 | 0.059 | 0.953 | 0.0008 | 0.010 | 0.084 | 0.933 | -0.0140 | 0.020 | -0.692 | 0.490 | -0.0072 | 0.024 | -0.302 | 0.763 | -0.0027 | 0.007 | -0.411 | 0.681 |
| mean FD x time | 0.0102 | 0.004 | 2.874 | 0.004** | 0.0056 | 0.004 | 1.375 | 0.170 | 0.0057 | 0.003 | 1.679 | 0.094. | 0.0072 | 0.004 | 1.622 | 0.106 | 0.0069 | 0.009 | 0.733 | 0.464 | 0.0068 | 0.011 | 0.615 | 0.539 | 0.0048 | 0.003 | 1.564 | 0.118 |
| tChol | 0.0007 | 0.003 | 0.266 | 0.790 | 0.0042 | 0.003 | 1.417 | 0.158 | 0.0047 | 0.003 | 1.670 | 0.096. | 0.0012 | 0.003 | 0.347 | 0.729 | 0.0059 | 0.008 | 0.790 | 0.430 | 0.0108 | 0.009 | 1.245 | 0.214 | 0.0044 | 0.002 | 1.789 | 0.075. |
| HDL | -0.0058 | 0.003 | -2.190 | 0.029* | -0.0071 | 0.003 | -2.436 | 0.016* | -0.0089 | 0.003 | -3.141 | 0.002** | -0.0016 | 0.003 | -0.493 | 0.622 | -0.0165 | 0.007 | -2.206 | 0.028* | -0.0226 | 0.009 | -2.614 | 0.010** | -0.0070 | 0.002 | -2.872 | 0.004** |
| sBP | -0.0027 | 0.003 | -0.943 | 0.347 | -0.0002 | 0.003 | -0.052 | 0.958 | -0.0028 | 0.003 | -0.901 | 0.368 | -0.0053 | 0.004 | -1.463 | 0.145 | 0.0142 | 0.008 | 1.717 | 0.087. | 0.0031 | 0.010 | 0.329 | 0.743 | -0.0015 | 0.003 | -0.556 | 0.579 |
| dBP | -0.0018 | 0.003 | -0.688 | 0.492 | -0.0029 | 0.003 | -0.966 | 0.335 | -0.0008 | 0.003 | -0.288 | 0.774 | -0.0001 | 0.003 | -0.040 | 0.968 | -0.0228 | 0.008 | -2.996 | 0.003** | -0.0128 | 0.009 | -1.456 | 0.147 | -0.0026 | 0.002 | -1.055 | 0.292 |
| age | 0.0037 | 0.002 | 1.654 | 0.099. | 0.0047 | 0.002 | 1.914 | 0.057. | 0.0024 | 0.002 | 0.990 | 0.323 | 0.0060 | 0.003 | 2.174 | 0.031* | -0.0049 | 0.006 | -0.783 | 0.434 | -0.0080 | 0.007 | -1.104 | 0.271 | 0.0042 | 0.002 | 2.061 | 0.040* |
| sex | -0.0031 | 0.005 | -0.569 | 0.570 | -0.0198 | 0.006 | -3.291 | 0.001** | -0.0102 | 0.006 | -1.744 | 0.083. | -0.0001 | 0.007 | -0.009 | 0.993 | -0.0198 | 0.015 | -1.286 | 0.200 | -0.0571 | 0.018 | -3.211 | 0.002** | -0.0078 | 0.005 | -1.542 | 0.125 |
| medication | -0.0104 | 0.005 | -2.056 | 0.041* | 0.0110 | 0.006 | 1.961 | 0.051. | 0.0040 | 0.005 | 0.743 | 0.458 | 0.0042 | 0.006 | 0.654 | 0.514 | -0.0038 | 0.014 | -0.263 | 0.793 | 0.0054 | 0.017 | 0.324 | 0.746 | 0.0039 | 0.005 | 0.834 | 0.405 |
| mean FD | -0.0005 | 0.002 | -0.288 | 0.774 | -0.0017 | 0.002 | -0.864 | 0.388 | 0.0049 | 0.002 | 2.698 | 0.007** | 0.0039 | 0.002 | 1.831 | 0.068. | 0.0020 | 0.005 | 0.410 | 0.682 | -0.0098 | 0.006 | -1.750 | 0.081. | 0.0017 | 0.002 | 1.038 | 0.300 |
| time | -0.0006 | 0.007 | -0.080 | 0.937 | -0.0063 | 0.008 | -0.755 | 0.451 | 0.0007 | 0.007 | 0.099 | 0.921 | -0.0025 | 0.009 | -0.268 | 0.789 | 0.0125 | 0.019 | 0.650 | 0.517 | -0.0093 | 0.023 | -0.406 | 0.685 | 0.0040 | 0.006 | 0.637 | 0.525 |
| R2m | 0.052 | | | | 0.089 | | | | 0.078 | | | | 0.036 | | | | 0.048 | | | | 0.074 | | | | 0.072 | | | |
| R2c | 0.463 | | | | 0.512 | | | | 0.555 | | | | 0.566 | | | | 0.520 | | | | 0.563 | | | | 0.521 | | | |

| Model B | DMN | | | | SAL/ VAN | | | | FPN | | | | LIM | | | | VIS | | | | SM | | | | global | | | |
|---|---|---|---|---|---|---|---|---|---|---|---|---|---|---|---|---|---|---|---|---|---|---|---|---|---|---|---|---|
| n of obs = 838 | | | | | | | | | | | | | | | | | | | | | | | | | | | | |
| n of part. = 240 | Est. | SE | t | p | Est. | SE | t | p | Est. | SE | t | p | Est. | SE | t | p | Est. | SE | t | p | Est. | SE | t | p | Est. | SE | t | p |
| LDL x time | -0.0092 | 0.004 | -2.301 | 0.023* | -0.0029 | 0.005 | -0.631 | 0.529 | -0.0047 | 0.004 | -1.257 | 0.211 | -0.0012 | 0.005 | -0.233 | 0.816 | -0.0037 | 0.011 | -0.346 | 0.730 | 0.0002 | 0.013 | 0.015 | 0.988 | -0.0037 | 0.003 | -1.104 | 0.270 |
| HDL x time | -0.0022 | 0.004 | -0.546 | 0.586 | 0.0002 | 0.005 | 0.040 | 0.968 | -0.0049 | 0.004 | -1.291 | 0.199 | -0.0100 | 0.005 | -1.921 | 0.056. | -0.0040 | 0.011 | -0.371 | 0.711 | -0.0204 | 0.013 | -1.581 | 0.116 | -0.0037 | 0.003 | -1.074 | 0.283 |
| sBP x time | 0.0005 | 0.005 | 0.107 | 0.915 | 0.0019 | 0.005 | 0.363 | 0.717 | -0.0001 | 0.004 | -0.032 | 0.974 | 0.0004 | 0.006 | 0.063 | 0.950 | -0.0060 | 0.012 | -0.486 | 0.627 | 0.0033 | 0.015 | 0.220 | 0.826 | 0.0007 | 0.004 | 0.182 | 0.856 |
| dBP x time | -0.0062 | 0.004 | -1.507 | 0.134 | -0.0117 | 0.005 | -2.478 | 0.014* | -0.0035 | 0.004 | -0.910 | 0.364 | -0.0054 | 0.005 | -1.011 | 0.313 | -0.0164 | 0.011 | -1.495 | 0.137 | -0.0244 | 0.013 | -1.848 | 0.067. | -0.0071 | 0.003 | -2.050 | 0.041* |
| age x time | -0.0020 | 0.003 | -0.594 | 0.554 | 0.0045 | 0.004 | 1.183 | 0.239 | -0.0031 | 0.003 | -1.000 | 0.319 | 0.0006 | 0.004 | 0.130 | 0.897 | -0.0061 | 0.009 | -0.705 | 0.482 | -0.0003 | 0.011 | -0.031 | 0.975 | -0.0001 | 0.003 | -0.043 | 0.966 |
| sex x time | -0.0007 | 0.008 | -0.084 | 0.933 | 0.0100 | 0.010 | 1.052 | 0.294 | 0.0072 | 0.008 | 0.936 | 0.351 | 0.0109 | 0.011 | 1.008 | 0.315 | -0.0036 | 0.022 | -0.165 | 0.869 | 0.0376 | 0.026 | 1.420 | 0.158 | 0.0053 | 0.007 | 0.765 | 0.445 |
| medication x time | -0.0051 | 0.008 | -0.633 | 0.528 | 0.0006 | 0.009 | 0.060 | 0.952 | -0.0006 | 0.008 | -0.080 | 0.936 | 0.0001 | 0.010 | 0.009 | 0.993 | -0.0057 | 0.021 | -0.266 | 0.790 | -0.0023 | 0.026 | -0.090 | 0.929 | -0.0013 | 0.007 | -0.188 | 0.851 |
| mean FD x time | 0.0092 | 0.004 | 2.566 | 0.011* | 0.0037 | 0.004 | 0.904 | 0.367 | 0.0047 | 0.003 | 1.405 | 0.161 | 0.0073 | 0.004 | 1.642 | 0.102 | 0.0059 | 0.010 | 0.623 | 0.534 | 0.0065 | 0.011 | 0.571 | 0.568 | 0.0034 | 0.003 | 1.125 | 0.261 |
| LDL | 0.0021 | 0.003 | 0.832 | 0.407 | 0.0040 | 0.003 | 1.412 | 0.159 | 0.0045 | 0.003 | 1.641 | 0.102 | 0.0009 | 0.003 | 0.268 | 0.789 | 0.0063 | 0.007 | 0.859 | 0.391 | 0.0076 | 0.009 | 0.890 | 0.375 | 0.0042 | 0.002 | 1.795 | 0.074. |
| HDL | -0.0054 | 0.003 | -2.166 | 0.031* | -0.0059 | 0.003 | -2.088 | 0.038* | -0.0076 | 0.003 | -2.831 | 0.005** | -0.0009 | 0.003 | -0.283 | 0.777 | -0.0140 | 0.007 | -1.942 | 0.053. | -0.0194 | 0.008 | -2.298 | 0.022* | -0.0056 | 0.002 | -2.425 | 0.016* |
| sBP | -0.0017 | 0.003 | -0.591 | 0.555 | 0.0010 | 0.003 | 0.308 | 0.759 | -0.0014 | 0.003 | -0.443 | 0.659 | -0.0048 | 0.004 | -1.267 | 0.206 | 0.0155 | 0.008 | 1.850 | 0.066. | 0.0020 | 0.010 | 0.201 | 0.841 | -0.0004 | 0.003 | -0.132 | 0.895 |
| dBP | -0.0027 | 0.003 | -1.014 | 0.312 | -0.0037 | 0.003 | -1.225 | 0.222 | -0.0015 | 0.003 | -0.508 | 0.612 | -0.0004 | 0.003 | -0.109 | 0.913 | -0.0236 | 0.008 | -3.055 | 0.003** | -0.0114 | 0.009 | -1.268 | 0.206 | -0.0032 | 0.002 | -1.299 | 0.195 |
| age | 0.0038 | 0.002 | 1.703 | 0.090. | 0.0051 | 0.002 | 2.055 | 0.041* | 0.0025 | 0.002 | 1.051 | 0.295 | 0.0059 | 0.003 | 2.090 | 0.038* | -0.0044 | 0.006 | -0.691 | 0.490 | -0.0080 | 0.007 | -1.076 | 0.283 | 0.0045 | 0.002 | 2.189 | 0.030* |
| sex | -0.0041 | 0.005 | -0.741 | 0.460 | -0.0193 | 0.006 | -3.150 | 0.002** | -0.0095 | 0.006 | -1.632 | 0.104 | -0.0007 | 0.007 | -0.098 | 0.922 | -0.0157 | 0.016 | -0.995 | 0.321 | -0.0530 | 0.018 | -2.884 | 0.004** | -0.0070 | 0.005 | -1.385 | 0.167 |
| medication | -0.0085 | 0.005 | -1.648 | 0.101 | 0.0121 | 0.006 | 2.090 | 0.038* | 0.0042 | 0.006 | 0.768 | 0.443 | 0.0037 | 0.007 | 0.551 | 0.582 | -0.0037 | 0.015 | -0.252 | 0.801 | 0.0054 | 0.017 | 0.310 | 0.757 | 0.0046 | 0.005 | 0.965 | 0.335 |
| mean FD | -0.0010 | 0.002 | -0.564 | 0.573 | -0.0020 | 0.002 | -1.018 | 0.309 | 0.0043 | 0.002 | 2.380 | 0.018* | 0.0047 | 0.002 | 2.165 | 0.031* | 0.0024 | 0.005 | 0.475 | 0.635 | -0.0100 | 0.006 | -1.731 | 0.084. | 0.0014 | 0.002 | 0.846 | 0.398 |
| time | 0.0020 | 0.008 | 0.262 | 0.794 | -0.0085 | 0.009 | -0.977 | 0.330 | 0.0015 | 0.007 | 0.215 | 0.830 | -0.0005 | 0.010 | -0.048 | 0.962 | 0.0118 | 0.020 | 0.587 | 0.558 | -0.0132 | 0.024 | -0.546 | 0.586 | 0.0041 | 0.006 | 0.642 | 0.521 |
| R2m | 0.051 | | | | 0.086 | | | | 0.066 | | | | 0.035 | | | | 0.043 | | | | 0.064 | | | | 0.065 | | | |
| R2c | 0.484 | | | | 0.531 | | | | 0.567 | | | | 0.586 | | | | 0.521 | | | | 0.561 | | | | 0.523 | | | |

Using linear mixed effects models, we test whether vascular risk factors (VRF) were associated with a change in resting-state functional connectivity (RSFC) within different functional networks that were defined by the MIST parcellation atlas (DMN, default mode network; SAL/VAN, salience and ventral attention network; FPN, fronto-parietal network; LIM, limbic network; VIS, visual network and SM, somatomotor network) and throughout the whole brain (global). Two separate models were conducted, where Model A included total cholesterol (tChol), high-density lipoprotein cholesterol (HDL), systolic blood pressure (sBP) and diastolic blood pressure (dBP) and Model B included low-density lipoprotein cholesterol (LDL), HDL, sBP and dBP, due to the severe multicollinearity between tChol and LDL. As random effects intercept and slope of RSFC for each individual were included in the models. Models were corrected for baseline age, sex and vascular medication (intake or non-intake of drugs against dyslipidemia and/or hypertension), for longitudinal mean frame-displacement (FD) and for the interactions of those covariates with time. Unstandardized estimates (Est.), standard errors (SE), t-values and p-values (***p<0.001, **p<0.01, *p<0.05) as well as marginal (R2m) and conditional (R2c) $R^2$ values are presented. Obs., observations; part, participants.

#### eTable 4. Association between VRF and longitudinal change in RSFC, using the Power parcellation

| Model A | DMN | | | | SAL | | | | VAN | | | | FPN | | | | LIM | | | | DAN | | | | VIS | | | | SM | | | | global | | | |
|---|---|---|---|---|---|---|---|---|---|---|---|---|---|---|---|---|---|---|---|---|---|---|---|---|---|---|---|---|---|---|---|---|---|---|---|---|
| n of obs. = 865 | | | | | | | | | | | | | | | | | | | | | | | | | | | | | | | | | | | | |
| n of part. = 247 | Est. | SE | t | p | Est. | SE | t | p | Est. | SE | t | p | Est. | SE | t | p | Est. | SE | t | p | Est. | SE | t | p | Est. | SE | t | p | Est. | SE | t | p | Est. | SE | t | p |
| tChol x time | -0.0085 | 0.003 | -2.536 | 0.012* | -0.0128 | 0.005 | -2.602 | 0.010* | -0.0117 | 0.008 | -1.542 | 0.125 | -0.0058 | 0.004 | -1.371 | 0.171 | 0.0083 | 0.011 | 0.753 | 0.453 | 0.0036 | 0.006 | 0.594 | 0.553 | -0.0128 | 0.009 | -1.350 | 0.179 | -0.0077 | 0.009 | -0.862 | 0.390 | -0.0047 | 0.003 | -1.602 | 0.110 |
| HDL x time | -0.0011 | 0.004 | -0.310 | 0.757 | -0.0028 | 0.005 | -0.539 | 0.591 | -0.0039 | 0.008 | -0.481 | 0.631 | -0.0056 | 0.005 | -1.232 | 0.219 | -0.0102 | 0.012 | -0.878 | 0.381 | -0.0065 | 0.007 | -0.994 | 0.321 | 0.0046 | 0.010 | 0.449 | 0.654 | -0.0119 | 0.010 | -1.245 | 0.215 | -0.0016 | 0.003 | -0.497 | 0.620 |
| sBP x time | 0.0001 | 0.004 | 0.022 | 0.983 | 0.0107 | 0.006 | 1.892 | 0.060. | 0.0010 | 0.009 | 0.114 | 0.910 | -0.0043 | 0.005 | -0.875 | 0.382 | 0.0026 | 0.013 | 0.206 | 0.837 | -0.0087 | 0.007 | -1.239 | 0.216 | 0.0034 | 0.011 | 0.311 | 0.756 | 0.0049 | 0.010 | 0.477 | 0.634 | 0.0014 | 0.003 | 0.418 | 0.676 |
| dBP x time | -0.0059 | 0.003 | -1.720 | 0.087. | -0.0035 | 0.005 | -0.690 | 0.491 | -0.0120 | 0.008 | -1.540 | 0.125 | -0.0002 | 0.004 | -0.049 | 0.961 | 0.0085 | 0.011 | 0.756 | 0.451 | -0.0027 | 0.006 | -0.426 | 0.670 | -0.0148 | 0.010 | -1.518 | 0.131 | -0.0146 | 0.009 | -1.582 | 0.116 | -0.0067 | 0.003 | -2.221 | 0.027* |
| age x time | 0.0003 | 0.003 | 0.124 | 0.902 | -0.0001 | 0.004 | -0.015 | 0.988 | 0.0042 | 0.006 | 0.676 | 0.500 | -0.0001 | 0.003 | -0.034 | 0.973 | -0.0012 | 0.009 | -0.136 | 0.892 | -0.0044 | 0.005 | -0.883 | 0.378 | -0.0072 | 0.008 | -0.936 | 0.351 | -0.0028 | 0.007 | -0.377 | 0.707 | -0.0009 | 0.002 | -0.363 | 0.717 |
| sex x time | 0.0033 | 0.007 | 0.484 | 0.629 | 0.0187 | 0.010 | 1.872 | 0.063. | -0.0012 | 0.015 | -0.080 | 0.936 | 0.0053 | 0.009 | 0.623 | 0.534 | 0.0175 | 0.022 | 0.785 | 0.434 | 0.0062 | 0.012 | 0.505 | 0.614 | 0.0012 | 0.019 | 0.063 | 0.950 | 0.0260 | 0.018 | 1.433 | 0.154 | 0.0052 | 0.006 | 0.875 | 0.382 |
| medication x time | -0.0002 | 0.006 | -0.030 | 0.976 | 0.0063 | 0.010 | 0.664 | 0.508 | -0.0252 | 0.015 | -1.730 | 0.085. | 0.0056 | 0.008 | 0.687 | 0.493 | 0.0224 | 0.021 | 1.060 | 0.291 | -0.0129 | 0.012 | -1.100 | 0.272 | -0.0133 | 0.018 | -0.729 | 0.467 | -0.0090 | 0.017 | -0.523 | 0.602 | -0.0031 | 0.006 | -0.548 | 0.584 |
| mean FD x time | 0.0088 | 0.003 | 2.932 | 0.004** | 0.0035 | 0.004 | 0.788 | 0.432 | 0.0115 | 0.007 | 1.711 | 0.088. | 0.0039 | 0.004 | 1.015 | 0.310 | -0.0047 | 0.009 | -0.502 | 0.616 | 0.0083 | 0.005 | 1.504 | 0.133 | 0.0094 | 0.009 | 1.102 | 0.271 | 0.0023 | 0.008 | 0.296 | 0.768 | 0.0047 | 0.003 | 1.761 | 0.079. |
| tChol | 0.0001 | 0.002 | 0.059 | 0.953 | 0.0014 | 0.003 | 0.427 | 0.669 | 0.0013 | 0.005 | 0.266 | 0.791 | 0.0016 | 0.003 | 0.482 | 0.630 | -0.0011 | 0.007 | -0.152 | 0.880 | 0.0129 | 0.004 | 2.908 | 0.004** | 0.0018 | 0.007 | 0.260 | 0.795 | 0.0046 | 0.006 | 0.764 | 0.446 | 0.0025 | 0.002 | 1.241 | 0.216 |
| HDL | -0.0071 | 0.002 | -3.288 | 0.001** | -0.0027 | 0.003 | -0.812 | 0.418 | -0.0124 | 0.005 | -2.463 | 0.015* | -0.0058 | 0.003 | -1.758 | 0.080. | 0.0097 | 0.007 | 1.375 | 0.171 | -0.0158 | 0.004 | -3.573 | 0.000*** | -0.0086 | 0.007 | -1.229 | 0.220 | -0.0107 | 0.006 | -1.771 | 0.078. | -0.0057 | 0.002 | -2.819 | 0.005** |
| sBP | -0.0006 | 0.002 | -0.254 | 0.800 | -0.0002 | 0.004 | -0.067 | 0.947 | -0.0013 | 0.006 | -0.240 | 0.811 | -0.0021 | 0.004 | -0.564 | 0.573 | 0.0010 | 0.008 | 0.122 | 0.903 | -0.0054 | 0.005 | -1.110 | 0.268 | 0.0158 | 0.008 | 2.046 | 0.042* | 0.0010 | 0.007 | 0.144 | 0.886 | 0.0001 | 0.002 | 0.062 | 0.950 |
| dBP | -0.0017 | 0.002 | -0.789 | 0.431 | -0.0021 | 0.003 | -0.635 | 0.526 | -0.0065 | 0.005 | -1.279 | 0.202 | 0.0058 | 0.003 | 1.712 | 0.088. | -0.0031 | 0.007 | -0.427 | 0.670 | -0.0051 | 0.004 | -1.138 | 0.256 | -0.0225 | 0.007 | -3.160 | 0.002** | -0.0081 | 0.006 | -1.319 | 0.189 | -0.0031 | 0.002 | -1.539 | 0.125 |
| age | 0.0033 | 0.002 | 1.808 | 0.072. | 0.0048 | 0.003 | 1.712 | 0.088. | 0.0012 | 0.004 | 0.294 | 0.769 | 0.0013 | 0.003 | 0.478 | 0.633 | 0.0112 | 0.006 | 1.886 | 0.061. | -0.0040 | 0.004 | -1.068 | 0.286 | -0.0065 | 0.006 | -1.109 | 0.268 | 0.0031 | 0.005 | 0.619 | 0.537 | 0.0037 | 0.002 | 2.195 | 0.029* |
| sex | 0.0071 | 0.004 | 1.612 | 0.108 | -0.0113 | 0.007 | -1.665 | 0.097. | -0.0170 | 0.010 | -1.644 | 0.101 | -0.0104 | 0.007 | -1.521 | 0.130 | 0.0263 | 0.015 | 1.803 | 0.073. | -0.0206 | 0.009 | -2.259 | 0.025* | -0.0108 | 0.014 | -0.747 | 0.456 | -0.0517 | 0.012 | -4.150 | 0.000*** | -0.0073 | 0.004 | -1.769 | 0.078. |
| medication | -0.0087 | 0.004 | -2.103 | 0.037* | -0.0009 | 0.006 | -0.137 | 0.892 | -0.0048 | 0.010 | -0.499 | 0.618 | 0.0001 | 0.006 | 0.021 | 0.984 | 0.0031 | 0.014 | 0.229 | 0.819 | 0.0100 | 0.008 | 1.187 | 0.236 | -0.0118 | 0.013 | -0.878 | 0.381 | 0.0017 | 0.012 | 0.143 | 0.886 | 0.0016 | 0.004 | 0.410 | 0.682 |
| mean FD | 0.0006 | 0.001 | 0.434 | 0.665 | 0.0057 | 0.002 | 2.550 | 0.011* | 0.0003 | 0.003 | 0.085 | 0.932 | 0.0019 | 0.002 | 0.916 | 0.360 | -0.0093 | 0.005 | -2.039 | 0.042* | 0.0035 | 0.003 | 1.214 | 0.225 | 0.0048 | 0.005 | 1.054 | 0.292 | 0.0004 | 0.004 | 0.103 | 0.918 | 0.0017 | 0.001 | 1.223 | 0.222 |
| time | -0.0057 | 0.006 | -0.932 | 0.353 | -0.0185 | 0.009 | -2.039 | 0.043* | 0.0069 | 0.014 | 0.495 | 0.621 | -0.0091 | 0.008 | -1.173 | 0.241 | -0.0277 | 0.020 | -1.376 | 0.170 | 0.0112 | 0.011 | 0.999 | 0.318 | 0.0170 | 0.017 | 0.972 | 0.332 | 0.0015 | 0.016 | 0.093 | 0.926 | 0.0026 | 0.005 | 0.481 | 0.631 |
| R2m | 0.065 | | | | 0.050 | | | | 0.048 | | | | 0.046 | | | | 0.082 | | | | 0.082 | | | | 0.041 | | | | 0.086 | | | | 0.079 | | | |
| R2c | 0.494 | | | | 0.516 | | | | 0.509 | | | | 0.565 | | | | 0.534 | | | | 0.534 | | | | 0.540 | | | | 0.547 | | | | 0.492 | | | |

| Model B | DMN | | | | SAL | | | | VAN | | | | FPN | | | | LIM | | | | DAN | | | | VIS | | | | SM | | | | global | | | |
|---|---|---|---|---|---|---|---|---|---|---|---|---|---|---|---|---|---|---|---|---|---|---|---|---|---|---|---|---|---|---|---|---|---|---|---|---|
| n of obs = 838 | | | | | | | | | | | | | | | | | | | | | | | | | | | | | | | | | | | | |
| n of part. = 240 | Est. | SE | t | p | Est. | SE | t | p | Est. | SE | t | p | Est. | SE | t | p | Est. | SE | t | p | Est. | SE | t | p | Est. | SE | t | p | Est. | SE | t | p | Est. | SE | t | p |
| LDL x time | -0.0060 | 0.003 | -1.744 | 0.083. | -0.0090 | 0.005 | -1.761 | 0.080. | -0.0090 | 0.008 | -1.167 | 0.245 | -0.0041 | 0.004 | -0.950 | 0.344 | 0.0083 | 0.011 | 0.748 | 0.455 | 0.0000 | 0.006 | 0.005 | 0.996 | -0.0089 | 0.010 | -0.923 | 0.357 | -0.0043 | 0.009 | -0.467 | 0.641 | -0.0030 | 0.003 | -1.020 | 0.309 |
| HDL x time | -0.0037 | 0.003 | -1.078 | 0.283 | -0.0063 | 0.005 | -1.220 | 0.224 | -0.0082 | 0.008 | -1.050 | 0.295 | -0.0078 | 0.004 | -1.778 | 0.077. | -0.0082 | 0.011 | -0.736 | 0.463 | -0.0051 | 0.006 | -0.822 | 0.411 | 0.0024 | 0.010 | 0.250 | 0.803 | -0.0150 | 0.009 | -1.610 | 0.109 | -0.0027 | 0.003 | -0.920 | 0.359 |
| sBP x time | -0.0007 | 0.004 | -0.180 | 0.857 | 0.0109 | 0.006 | 1.847 | 0.066. | 0.0022 | 0.009 | 0.245 | 0.807 | -0.0042 | 0.005 | -0.832 | 0.407 | 0.0040 | 0.013 | 0.309 | 0.758 | -0.0076 | 0.007 | -1.066 | 0.287 | 0.0050 | 0.011 | 0.441 | 0.660 | 0.0058 | 0.011 | 0.537 | 0.592 | 0.0019 | 0.003 | 0.559 | 0.577 |
| dBP x time | -0.0054 | 0.004 | -1.525 | 0.129 | -0.0038 | 0.005 | -0.722 | 0.471 | -0.0119 | 0.008 | -1.493 | 0.137 | 0.0008 | 0.004 | 0.176 | 0.861 | 0.0076 | 0.011 | 0.662 | 0.509 | 0.0003 | 0.006 | 0.049 | 0.961 | -0.0177 | 0.010 | -1.776 | 0.078. | -0.0154 | 0.010 | -1.609 | 0.110 | -0.0071 | 0.003 | -2.366 | 0.019* |
| age x time | 0.0004 | 0.003 | 0.137 | 0.891 | 0.0003 | 0.004 | 0.079 | 0.937 | 0.0037 | 0.006 | 0.579 | 0.564 | -0.0002 | 0.004 | -0.051 | 0.959 | 0.0008 | 0.009 | 0.090 | 0.928 | -0.0044 | 0.005 | -0.871 | 0.384 | -0.0071 | 0.008 | -0.890 | 0.375 | -0.0029 | 0.008 | -0.384 | 0.701 | -0.0005 | 0.002 | -0.214 | 0.831 |
| sex x time | 0.0011 | 0.007 | 0.152 | 0.879 | 0.0174 | 0.011 | 1.652 | 0.101 | -0.0034 | 0.016 | -0.213 | 0.831 | 0.0063 | 0.009 | 0.706 | 0.481 | 0.0147 | 0.023 | 0.636 | 0.526 | 0.0076 | 0.013 | 0.606 | 0.545 | -0.0019 | 0.020 | -0.096 | 0.924 | 0.0249 | 0.019 | 1.300 | 0.196 | 0.0048 | 0.006 | 0.799 | 0.426 |
| medication x time | 0.0024 | 0.007 | 0.346 | 0.730 | 0.0085 | 0.010 | 0.830 | 0.408 | -0.0274 | 0.016 | -1.761 | 0.080. | 0.0041 | 0.009 | 0.477 | 0.634 | 0.0260 | 0.022 | 1.160 | 0.248 | -0.0196 | 0.012 | -1.586 | 0.113 | -0.0077 | 0.019 | -0.396 | 0.693 | -0.0092 | 0.019 | -0.492 | 0.624 | -0.0012 | 0.006 | -0.212 | 0.832 |
| mean FD x time | 0.0071 | 0.003 | 2.367 | 0.019* | 0.0016 | 0.004 | 0.352 | 0.725 | 0.0104 | 0.007 | 1.516 | 0.131 | 0.0029 | 0.004 | 0.754 | 0.451 | -0.0055 | 0.010 | -0.575 | 0.566 | 0.0076 | 0.006 | 1.382 | 0.168 | 0.0081 | 0.009 | 0.939 | 0.349 | 0.0027 | 0.008 | 0.329 | 0.742 | 0.0034 | 0.003 | 1.298 | 0.196 |
| LDL | 0.0012 | 0.002 | 0.552 | 0.581 | 0.0026 | 0.003 | 0.811 | 0.418 | 0.0035 | 0.005 | 0.714 | 0.476 | 0.0023 | 0.003 | 0.698 | 0.486 | -0.0021 | 0.007 | -0.311 | 0.756 | 0.0147 | 0.004 | 3.405 | 0.001*** | 0.0027 | 0.007 | 0.394 | 0.694 | 0.0020 | 0.006 | 0.325 | 0.746 | 0.0029 | 0.002 | 1.518 | 0.130 |
| HDL | -0.0068 | 0.002 | -3.295 | 0.001** | -0.0022 | 0.003 | -0.703 | 0.483 | -0.0126 | 0.005 | -2.604 | 0.010** | -0.0057 | 0.003 | -1.796 | 0.074. | 0.0142 | 0.007 | 2.106 | 0.036* | -0.0135 | 0.004 | -3.171 | 0.002** | -0.0075 | 0.007 | -1.100 | 0.272 | -0.0102 | 0.006 | -1.717 | 0.087. | -0.0048 | 0.002 | -2.545 | 0.012* |
| sBP | -0.0002 | 0.002 | -0.100 | 0.920 | 0.0005 | 0.004 | 0.141 | 0.888 | -0.0007 | 0.006 | -0.121 | 0.904 | -0.0014 | 0.004 | -0.375 | 0.708 | 0.0015 | 0.008 | 0.189 | 0.850 | -0.0028 | 0.005 | -0.576 | 0.565 | 0.0163 | 0.008 | 2.050 | 0.042* | 0.0018 | 0.007 | 0.255 | 0.799 | 0.0009 | 0.002 | 0.429 | 0.668 |
| dBP | -0.0021 | 0.002 | -0.932 | 0.352 | -0.0028 | 0.003 | -0.839 | 0.402 | -0.0075 | 0.005 | -1.462 | 0.148 | 0.0057 | 0.003 | 1.666 | 0.097. | -0.0027 | 0.007 | -0.376 | 0.708 | -0.0067 | 0.005 | -1.462 | 0.145 | -0.0229 | 0.007 | -3.140 | 0.002** | -0.0079 | 0.006 | -1.239 | 0.217 | -0.0037 | 0.002 | -1.834 | 0.068. |
| age | 0.0033 | 0.002 | 1.832 | 0.068. | 0.0047 | 0.003 | 1.707 | 0.089. | 0.0017 | 0.004 | 0.409 | 0.683 | 0.0011 | 0.003 | 0.374 | 0.709 | 0.0098 | 0.006 | 1.662 | 0.098. | -0.0030 | 0.004 | -0.798 | 0.426 | -0.0062 | 0.006 | -1.031 | 0.303 | 0.0037 | 0.005 | 0.709 | 0.479 | 0.0039 | 0.002 | 2.352 | 0.020* |
| sex | 0.0062 | 0.005 | 1.373 | 0.171 | -0.0132 | 0.007 | -1.918 | 0.056. | -0.0203 | 0.011 | -1.930 | 0.055. | -0.0115 | 0.007 | -1.646 | 0.101 | 0.0331 | 0.015 | 2.256 | 0.025* | -0.0219 | 0.009 | -2.356 | 0.019* | -0.0085 | 0.015 | -0.574 | 0.567 | -0.0520 | 0.013 | -4.021 | 0.000*** | -0.0070 | 0.004 | -1.681 | 0.094. |
| medication | -0.0071 | 0.004 | -1.665 | 0.097. | 0.0011 | 0.006 | 0.168 | 0.867 | -0.0005 | 0.010 | -0.049 | 0.961 | 0.0003 | 0.007 | 0.041 | 0.967 | 0.0016 | 0.014 | 0.113 | 0.910 | 0.0130 | 0.009 | 1.482 | 0.140 | -0.0111 | 0.014 | -0.792 | 0.429 | 0.0001 | 0.012 | 0.010 | 0.992 | 0.0027 | 0.004 | 0.693 | 0.489 |
| mean FD | 0.0002 | 0.001 | 0.108 | 0.914 | 0.0057 | 0.002 | 2.567 | 0.011* | -0.0003 | 0.003 | -0.094 | 0.925 | 0.0014 | 0.002 | 0.666 | 0.505 | -0.0096 | 0.005 | -2.102 | 0.036* | 0.0033 | 0.003 | 1.118 | 0.264 | 0.0049 | 0.005 | 1.064 | 0.288 | 0.0008 | 0.004 | 0.203 | 0.839 | 0.0014 | 0.001 | 1.022 | 0.307 |
| time | -0.0048 | 0.006 | -0.749 | 0.455 | -0.0189 | 0.010 | -1.967 | 0.051. | 0.0099 | 0.015 | 0.675 | 0.500 | -0.0094 | 0.008 | -1.148 | 0.253 | -0.0268 | 0.021 | -1.282 | 0.202 | 0.0125 | 0.012 | 1.077 | 0.282 | 0.0166 | 0.018 | 0.911 | 0.364 | 0.0024 | 0.017 | 0.138 | 0.891 | 0.0020 | 0.005 | 0.361 | 0.718 |
| R2m | 0.058 | | | | 0.050 | | | | 0.050 | | | | 0.044 | | | | 0.057 | | | | 0.082 | | | | 0.038 | | | | 0.082 | | | | 0.074 | | | |
| R2c | 0.519 | | | | 0.523 | | | | 0.509 | | | | 0.579 | | | | 0.592 | | | | 0.536 | | | | 0.548 | | | | 0.551 | | | | 0.490 | | | |

Using linear mixed effects models, we test whether vascular risk factors (VRF) were associated with a change in resting-state functional connectivity (RSFC) within different functional networks that were defined by the Power parcellation atlas (DMN, default mode network; SAL, salience network; VAN, ventral attention network; FPN, fronto-parietal network; LIM, limbic network; DAN, dorsal attention network; VIS, visual network and SM, somatomotor network) and throughout the whole brain (global). Two separate models were conducted, where Model A included total cholesterol (tChol), high-density lipoprotein cholesterol (HDL), systolic blood pressure (sBP) and diastolic blood pressure (dBP) and Model B included low-density lipoprotein cholesterol (LDL), HDL, sBP and dBP, due to the severe multicollinearity between tChol and LDL. As random effects intercept and slope of RSFC for each individual were included in the models. Models were corrected for baseline age, sex and vascular medication (intake or non-intake of drugs against dyslipidemia and/or hypertension), for longitudinal mean frame-displacement (FD) and for the interactions of those covariates with time. Unstandardized estimates (Est.), standard errors (SE), t-values and p-values (***$p<0.001$, **$p<0.01$, *$p<0.05$) as well as marginal (R2m) and conditional (R2c) $R^2$ values are presented. Obs., observations; part, participants.

#### eTable 5. Association between AD biomarkers and longitudinal changes in RSFC, using the Schaefer parcellation

| Aβ<br>n of obs. = 340<br>n of part. = 91 | DMN | | | | SAL/ VAN | | | | FPN | | | | LIM | | | |
|---|---|---|---|---|---|---|---|---|---|---|---|---|---|---|---|---|
| | **Est.** | **SE** | **t** | **p** | **Est.** | **SE** | **t** | **p** | **Est.** | **SE** | **t** | **p** | **Est.** | **SE** | **t** | **p** |
| Aβ x time | -0.0008 | 0.002 | -0.359 | 0.721 | 0.0006 | 0.003 | 0.231 | 0.818 | 0.0014 | 0.002 | 0.596 | 0.552 | -0.0018 | 0.004 | -0.468 | 0.642 |
| age x time | -0.0005 | 0.002 | -0.208 | 0.836 | 0.0005 | 0.003 | 0.168 | 0.867 | -0.0025 | 0.002 | -1.054 | 0.293 | -0.0009 | 0.004 | -0.241 | 0.810 |
| sex x time | 0.0028 | 0.005 | 0.551 | 0.584 | 0.0000 | 0.006 | -0.001 | 0.999 | 0.0023 | 0.005 | 0.434 | 0.665 | 0.0079 | 0.008 | 0.967 | 0.338 |
| mean FD x time | 0.0026 | 0.002 | 1.169 | 0.245 | 0.0014 | 0.003 | 0.538 | 0.591 | 0.0025 | 0.002 | 1.070 | 0.286 | 0.0060 | 0.004 | 1.635 | 0.106 |
| Aβ | -0.0037 | 0.004 | -0.877 | 0.383 | -0.0037 | 0.005 | -0.780 | 0.437 | -0.0014 | 0.005 | -0.254 | 0.800 | 0.0046 | 0.009 | 0.534 | 0.595 |
| age | -0.0004 | 0.004 | -0.088 | 0.930 | -0.0006 | 0.005 | -0.122 | 0.903 | 0.0004 | 0.005 | 0.077 | 0.939 | -0.0041 | 0.009 | -0.466 | 0.643 |
| sex | -0.0108 | 0.010 | -1.124 | 0.265 | -0.0253 | 0.011 | -2.302 | 0.024* | -0.0090 | 0.012 | -0.726 | 0.470 | -0.0122 | 0.020 | -0.612 | 0.542 |
| mean FD | -0.0007 | 0.003 | -0.244 | 0.808 | 0.0032 | 0.004 | 0.883 | 0.378 | 0.0065 | 0.004 | 1.834 | 0.068. | 0.0212 | 0.006 | 3.825 | 0.000*** |
| time | -0.0019 | 0.004 | -0.424 | 0.674 | -0.0041 | 0.005 | -0.824 | 0.411 | -0.0020 | 0.005 | -0.447 | 0.655 | 0.0062 | 0.007 | 0.878 | 0.384 |
| R2m | 0.018 | | | | 0.048 | | | | 0.020 | | | | 0.072 | | | |
| R2c | 0.540 | | | | 0.502 | | | | 0.617 | | | | 0.668 | | | |

| Aβ<br>n of obs. = 340<br>n of part. = 91 | DAN | | | | VIS | | | | SM | | | | global | | | |
|---|---|---|---|---|---|---|---|---|---|---|---|---|---|---|---|---|
| | **Est.** | **SE** | **t** | **p** | **Est.** | **SE** | **t** | **p** | **Est.** | **SE** | **t** | **p** | **Est.** | **SE** | **t** | **p** |
| Aβ x time | 0.0047 | 0.004 | 1.320 | 0.188 | 0.0001 | 0.004 | 0.023 | 0.982 | -0.0027 | 0.004 | -0.626 | 0.532 | 0.0003 | 0.002 | 0,128 | 0,898 |
| age x time | -0.0007 | 0.004 | -0.209 | 0.835 | -0.0012 | 0.004 | -0.293 | 0.770 | 0.0022 | 0.004 | 0.517 | 0.606 | -0.0008 | 0.002 | -0,401 | 0,689 |
| sex x time | 0.0062 | 0.008 | 0.806 | 0.421 | -0.0058 | 0.009 | -0.680 | 0.498 | 0.0100 | 0.009 | 1.102 | 0.272 | 0.0021 | 0.005 | 0,464 | 0,643 |
| mean FD x time | 0.0005 | 0.003 | 0.142 | 0.887 | 0.0046 | 0.004 | 1.224 | 0.222 | 0.0021 | 0.004 | 0.523 | 0.602 | 0.0024 | 0.002 | 1,154 | 0,250 |
| Aβ | -0.0055 | 0.007 | -0.761 | 0.449 | -0.0073 | 0.008 | -0.972 | 0.334 | -0.0052 | 0.008 | -0.636 | 0.527 | -0.0017 | 0.004 | -0,428 | 0,670 |
| age | 0.0004 | 0.007 | 0.057 | 0.955 | -0.0089 | 0.008 | -1.154 | 0.252 | -0.0004 | 0.008 | -0.051 | 0.959 | -0.0017 | 0.004 | -0,410 | 0,683 |
| sex | -0.0423 | 0.017 | -2.535 | 0.013* | -0.0367 | 0.017 | -2.107 | 0.038* | -0.0479 | 0.019 | -2.534 | 0.013* | -0.0198 | 0.009 | -2,119 | 0,037* |
| mean FD | 0.0129 | 0.005 | 2.492 | 0.013* | 0.0067 | 0.005 | 1.237 | 0.217 | 0.0019 | 0.006 | 0.322 | 0.747 | 0.0053 | 0.003 | 1,801 | 0,073. |
| time | 0.0003 | 0.007 | 0.050 | 0.960 | 0.0007 | 0.007 | 0.096 | 0.923 | -0.0034 | 0.008 | -0.431 | 0.667 | 0.0005 | 0.004 | 0,119 | 0,906 |
| R2m | 0.083 | | | | 0.053 | | | | 0.053 | | | | 0.049 | | | |
| R2c | 0.579 | | | | 0.550 | | | | 0.557 | | | | 0.537 | | | |

| *Tau*<br>n of obs = 340<br>n of part. = 91 | DMN | | | | SAL/ VAN | | | | FPN | | | | LIM | | | |
|---|---|---|---|---|---|---|---|---|---|---|---|---|---|---|---|---|
| | **Est.** | **SE** | **t** | **p** | **Est.** | **SE** | **t** | **p** | **Est.** | **SE** | **t** | **p** | **Est.** | **SE** | **t** | **p** |
| tau x time | -0.0021 | 0.002 | -0.876 | 0.384 | -0.0007 | 0.003 | -0.236 | 0.814 | 0.0002 | 0.002 | 0.084 | 0.933 | -0.0067 | 0.004 | -1.785 | 0.079. |
| age x time | -0.0001 | 0.002 | -0.033 | 0.974 | 0.0008 | 0.003 | 0.288 | 0.774 | -0.0022 | 0.002 | -0.926 | 0.356 | 0.0006 | 0.004 | 0.175 | 0.862 |
| sex x time | 0.0029 | 0.005 | 0.571 | 0.570 | 0.0002 | 0.006 | 0.026 | 0.979 | 0.0023 | 0.005 | 0.441 | 0.660 | 0.0078 | 0.008 | 0.975 | 0.334 |
| mean FD x time | 0.0027 | 0.002 | 1.182 | 0.240 | 0.0014 | 0.003 | 0.538 | 0.591 | 0.0025 | 0.002 | 1.057 | 0.291 | 0.0058 | 0.004 | 1.619 | 0.109 |
| tau | -0.0002 | 0.004 | -0.038 | 0.970 | -0.0029 | 0.005 | -0.589 | 0.557 | -0.0001 | 0.006 | -0.017 | 0.986 | -0.0019 | 0.009 | -0.221 | 0.825 |
| age | -0.0010 | 0.004 | -0.231 | 0.818 | -0.0005 | 0.005 | -0.110 | 0.913 | 0.0002 | 0.006 | 0.032 | 0.975 | -0.0028 | 0.009 | -0.311 | 0.756 |
| sex | -0.0097 | 0.010 | -1.004 | 0.318 | -0.0242 | 0.011 | -2.198 | 0.031* | -0.0089 | 0.012 | -0.719 | 0.474 | -0.0118 | 0.020 | -0.595 | 0.553 |
| mean FD | -0.0007 | 0.003 | -0.244 | 0.807 | 0.0032 | 0.004 | 0.891 | 0.374 | 0.0065 | 0.004 | 1.837 | 0.067· | 0.0210 | 0.006 | 3.789 | 0.000*** |
| time | -0.0020 | 0.004 | -0.441 | 0.661 | -0.0043 | 0.005 | -0.853 | 0.394 | -0.0021 | 0.005 | -0.465 | 0.642 | 0.0063 | 0.007 | 0.904 | 0.370 |
| R2m | 0.013 | | | | 0.044 | | | | 0.019 | | | | 0.074 | | | |
| R2c | 0.542 | | | | 0.502 | | | | 0.617 | | | | 0.666 | | | |

| *Tau*<br>n of obs = 340<br>n of part. = 91 | DAN | | | | VIS | | | | SM | | | | global | | | |
|---|---|---|---|---|---|---|---|---|---|---|---|---|---|---|---|---|
| | **Est.** | **SE** | **t** | **p** | **Est.** | **SE** | **t** | **p** | **Est.** | **SE** | **t** | **p** | **Est.** | **SE** | **t** | **p** |
| tau x time | 0.0021 | 0.004 | 0.590 | 0.556 | -0.0008 | 0.004 | -0.193 | 0.847 | 0.0009 | 0.004 | 0.204 | 0.839 | -0.0002 | 0.002 | -0,072 | 0,942 |
| age x time | -0.0002 | 0.004 | -0.056 | 0.955 | -0.0009 | 0.004 | -0.222 | 0.825 | 0.0013 | 0.004 | 0.305 | 0.761 | -0.0007 | 0.002 | -0,342 | 0,732 |
| sex x time | 0.0067 | 0.008 | 0.872 | 0.384 | -0.0055 | 0.009 | -0.641 | 0.522 | 0.0103 | 0.009 | 1.133 | 0.259 | 0.0022 | 0.005 | 0,484 | 0,629 |
| mean FD x time | 0.0004 | 0.003 | 0.104 | 0.918 | 0.0047 | 0.004 | 1.235 | 0.218 | 0.0022 | 0.004 | 0.551 | 0.582 | 0.0024 | 0.002 | 1,156 | 0,249 |
| tau | -0.0043 | 0.007 | -0.576 | 0.566 | -0.0081 | 0.008 | -1.049 | 0.297 | -0.0089 | 0.008 | -1.069 | 0.288 | -0.0017 | 0.004 | -0,405 | 0,686 |
| age | 0.0004 | 0.008 | 0.051 | 0.960 | -0.0081 | 0.008 | -1.043 | 0.300 | 0.0009 | 0.008 | 0.111 | 0.912 | -0.0016 | 0.004 | -0,378 | 0,706 |
| sex | -0.0418 | 0.017 | -2.497 | 0.015* | -0.0341 | 0.017 | -1.965 | 0.053. | -0.0456 | 0.019 | -2.429 | 0.017* | -0.0193 | 0.009 | -2,068 | 0,042* |
| mean FD | 0.0131 | 0.005 | 2.529 | 0.012* | 0.0068 | 0.005 | 1.247 | 0.213 | 0.0020 | 0.006 | 0.337 | 0.736 | 0.0053 | 0.003 | 1,806 | 0,072. |
| time | -0.0003 | 0.007 | -0.040 | 0.968 | 0.0005 | 0.007 | 0.063 | 0.950 | -0.0035 | 0.008 | -0.441 | 0.659 | 0.0004 | 0.004 | 0,099 | 0,921 |
| R2m | 0.077 | | | | 0.051 | | | | 0.054 | | | | 0.048 | | | |
| R2c | 0.579 | | | | 0.549 | | | | 0.555 | | | | 0.536 | | | |

Using linear mixed effects models, we test whether global amyloid-β (Aβ) and entorhinal *tau* deposition were associated with a change in resting-state functional connectivity (RSFC) within different functional networks that were defined by the Schaefer parcellation atlas (DMN, default mode network; SAL/VAN, salience and ventral attention network; FPN, fronto-parietal network; LIM, limbic network; DAN, dorsal attention network; VIS, visual network and SM, somatomotor network) and throughout the whole brain (global). As random effects intercept and slope of RSFC for each individual were included in the models. Models were corrected for baseline age and sex, for longitudinal mean frame-displacement (FD) and for the interactions of those covariates with time. Unstandardized estimates (Est.), standard errors (SE), t-values and p-values (***$p<0.001$, *$p<0.05$) as well as marginal (R2m) and conditional (R2c) $R^2$ values are presented. Obs., observations; part, participants.

#### eTable 6. Association between AD biomarkers and longitudinal changes in RSFC, using the MIST parcellation

| Aβ | DMN | | | | SAL/ VAN | | | | FPN | | | | LIM | | | | VIS | | | | SM | | | | global | | | |
|---|---|---|---|---|---|---|---|---|---|---|---|---|---|---|---|---|---|---|---|---|---|---|---|---|---|---|---|---|
| n of obs. = 340 | | | | | | | | | | | | | | | | | | | | | | | | | | | | |
| n of part. = 91 | Est. | SE | t | p | Est. | SE | t | p | Est. | SE | t | p | Est. | SE | t | p | Est. | SE | t | p | Est. | SE | t | p | Est. | SE | t | p |
| Aβ x time | -0.0007 | 0.002 | -0.295 | 0.769 | 0.0014 | 0.003 | 0.534 | 0.594 | 0.0026 | 0.002 | 1.169 | 0.244 | -0.0025 | 0.002 | -1.064 | 0.291 | -0.0015 | 0.005 | -0.291 | 0.772 | -0.0068 | 0.006 | -1.059 | 0.291 | 0.0002 | 0.002 | 0.111 | 0.912 |
| age x time | -0.0003 | 0.002 | -0.129 | 0.897 | 0.0006 | 0.003 | 0.233 | 0.816 | -0.0023 | 0.002 | -1.047 | 0.296 | -0.0011 | 0.002 | -0.479 | 0.634 | 0.0002 | 0.005 | 0.034 | 0.973 | 0.0034 | 0.006 | 0.541 | 0.589 | -0.0006 | 0.002 | -0.313 | 0.755 |
| sex x time | 0.0058 | 0.005 | 1.166 | 0.249 | 0.0039 | 0.005 | 0.708 | 0.480 | 0.0018 | 0.005 | 0.385 | 0.701 | 0.0035 | 0.005 | 0.692 | 0.492 | -0.0048 | 0.011 | -0.427 | 0.670 | 0.0085 | 0.014 | 0.615 | 0.539 | 0.0018 | 0.004 | 0.434 | 0.665 |
| mean FD x time | 0.0021 | 0.002 | 0.937 | 0.351 | 0.0008 | 0.002 | 0.314 | 0.754 | 0.0023 | 0.002 | 1.092 | 0.276 | 0.0032 | 0.002 | 1.414 | 0.161 | 0.0072 | 0.005 | 1.414 | 0.159 | 0.0020 | 0.006 | 0.313 | 0.755 | 0.0024 | 0.002 | 1.309 | 0.192 |
| Aβ | -0.0020 | 0.004 | -0.493 | 0.624 | -0.0051 | 0.004 | -1.137 | 0.259 | -0.0035 | 0.005 | -0.757 | 0.451 | 0.0032 | 0.004 | 0.740 | 0.461 | -0.0129 | 0.010 | -1.259 | 0.211 | -0.0139 | 0.013 | -1.106 | 0.272 | -0.0019 | 0.004 | -0.507 | 0.613 |
| age | -0.0027 | 0.004 | -0.645 | 0.521 | 0.0014 | 0.005 | 0.313 | 0.755 | 0.0007 | 0.005 | 0.144 | 0.886 | -0.0021 | 0.004 | -0.469 | 0.640 | -0.0064 | 0.010 | -0.612 | 0.542 | -0.0029 | 0.013 | -0.230 | 0.819 | -0.0009 | 0.004 | -0.242 | 0.809 |
| sex | -0.0101 | 0.009 | -1.073 | 0.287 | -0.0281 | 0.010 | -2.717 | 0.008** | -0.0085 | 0.011 | -0.804 | 0.424 | 0.0061 | 0.010 | 0.605 | 0.547 | -0.0422 | 0.024 | -1.786 | 0.078. | -0.0647 | 0.029 | -2.236 | 0.028* | -0.0127 | 0.009 | -1.483 | 0.142 |
| mean FD | -0.0011 | 0.003 | -0.355 | 0.723 | 0.0058 | 0.003 | 1.701 | 0.090. | 0.0076 | 0.003 | 2.401 | 0.017* | 0.0077 | 0.003 | 2.465 | 0.014* | 0.0077 | 0.007 | 1.046 | 0.296 | 0.0015 | 0.009 | 0.159 | 0.873 | 0.0051 | 0.003 | 1.895 | 0.059, |
| time | -0.0052 | 0.004 | -1.230 | 0.225 | -0.0032 | 0.005 | -0.683 | 0.495 | -0.0010 | 0.004 | -0.250 | 0.802 | 0.0003 | 0.004 | 0.070 | 0.945 | 0.0022 | 0.010 | 0.228 | 0.820 | 0.0024 | 0.012 | 0.203 | 0.839 | 0.0007 | 0.004 | 0.206 | 0.837 |
| R2m | 0.018 | | | | 0.072 | | | | 0.036 | | | | 0.037 | | | | 0.042 | | | | 0.048 | | | | 0.036 | | | |
| R2c | 0.497 | | | | 0.501 | | | | 0.585 | | | | 0.575 | | | | 0.544 | | | | 0.541 | | | | 0.531 | | | |

| *Tau* | DMN | | | | SAL/ VAN | | | | FPN | | | | LIM | | | | VIS | | | | SM | | | | global | | | |
|---|---|---|---|---|---|---|---|---|---|---|---|---|---|---|---|---|---|---|---|---|---|---|---|---|---|---|---|---|
| n of obs. = 340 | | | | | | | | | | | | | | | | | | | | | | | | | | | | |
| n of part. = 91 | Est. | SE | t | p | Est. | SE | t | p | Est. | SE | t | p | Est. | SE | t | p | Est. | SE | t | p | Est. | SE | t | p | Est. | SE | t | p |
| tau x time | -0.0017 | 0.002 | -0.718 | 0.476 | 0.0024 | 0.003 | 0.897 | 0.371 | 0.0006 | 0.002 | 0.273 | 0.785 | -0.0026 | 0.002 | -1.073 | 0.287 | -0.0015 | 0.005 | -0.280 | 0.780 | -0.0047 | 0.007 | -0.720 | 0.472 | -0.0002 | 0.002 | -0.100 | 0.920 |
| age x time | 0.0000 | 0.002 | 0.012 | 0.990 | 0.0002 | 0.003 | 0.080 | 0.936 | -0.0018 | 0.002 | -0.843 | 0.400 | -0.0010 | 0.002 | -0.416 | 0.679 | 0.0003 | 0.005 | 0.052 | 0.959 | 0.0031 | 0.006 | 0.484 | 0.629 | -0.0005 | 0.002 | -0.252 | 0.802 |
| sex x time | 0.0058 | 0.005 | 1.174 | 0.246 | 0.0039 | 0.006 | 0.712 | 0.477 | 0.0019 | 0.005 | 0.407 | 0.685 | 0.0034 | 0.005 | 0.661 | 0.511 | -0.0042 | 0.011 | -0.368 | 0.713 | 0.0097 | 0.014 | 0.703 | 0.483 | 0.0019 | 0.004 | 0.453 | 0.651 |
| mean FD x time | 0.0021 | 0.002 | 0.950 | 0.345 | 0.0008 | 0.002 | 0.318 | 0.751 | 0.0023 | 0.002 | 1.065 | 0.288 | 0.0032 | 0.002 | 1.438 | 0.154 | 0.0073 | 0.005 | 1.434 | 0.153 | 0.0023 | 0.006 | 0.363 | 0.717 | 0.0024 | 0.002 | 1.311 | 0.191 |
| tau | -0.0006 | 0.004 | -0.136 | 0.892 | -0.0024 | 0.005 | -0.520 | 0.605 | -0.0005 | 0.005 | -0.103 | 0.919 | 0.0029 | 0.004 | 0.645 | 0.520 | -0.0141 | 0.010 | -1.353 | 0.180 | -0.0301 | 0.012 | -2.420 | 0.018* | -0.0012 | 0.004 | -0.314 | 0.754 |
| age | -0.0029 | 0.004 | -0.685 | 0.495 | 0.0011 | 0.005 | 0.229 | 0.820 | 0.0001 | 0.005 | 0.026 | 0.979 | -0.0022 | 0.005 | -0.482 | 0.631 | -0.0051 | 0.011 | -0.482 | 0.631 | 0.0023 | 0.013 | 0.183 | 0.855 | -0.0010 | 0.004 | -0.249 | 0.804 |
| sex | -0.0093 | 0.009 | -0.993 | 0.324 | -0.0273 | 0.010 | -2.632 | 0.010* | -0.0081 | 0.011 | -0.764 | 0.447 | 0.0058 | 0.010 | 0.582 | 0.562 | -0.0375 | 0.024 | -1.593 | 0.115 | -0.0572 | 0.028 | -2.036 | 0.045* | -0.0121 | 0.009 | -1.423 | 0.158 |
| mean FD | -0.0011 | 0.003 | -0.357 | 0.721 | 0.0060 | 0.003 | 1.752 | 0.081. | 0.0077 | 0.003 | 2.417 | 0.016* | 0.0076 | 0.003 | 2.435 | 0.016* | 0.0078 | 0.007 | 1.064 | 0.288 | 0.0015 | 0.009 | 0.169 | 0.866 | 0.0051 | 0.003 | 1.901 | 0.058, |
| time | -0.0053 | 0.004 | -1.233 | 0.223 | -0.0032 | 0.005 | -0.687 | 0.493 | -0.0012 | 0.004 | -0.295 | 0.769 | 0.0005 | 0.004 | 0.103 | 0.918 | 0.0018 | 0.010 | 0.182 | 0.856 | 0.0016 | 0.012 | 0.137 | 0.891 | 0.0007 | 0.004 | 0.187 | 0.852 |
| R2m | 0.017 | | | | 0.064 | | | | 0.029 | | | | 0.035 | | | | 0.041 | | | | 0.073 | | | | 0.034 | | | |
| R2c | 0.499 | | | | 0.502 | | | | 0.585 | | | | 0.575 | | | | 0.544 | | | | 0.536 | | | | 0.531 | | | |

Using linear mixed effects models, we test whether global amyloid-β (Aβ) and entorhinal *tau* deposition were associated with a change in resting-state functional connectivity (RSFC) within different functional networks that were defined by the MIST parcellation atlas (DMN, default mode network; SAL/VAN, salience and ventral attention network; FPN, fronto-parietal network; LIM, limbic network; VIS, visual network and SM, somatomotor network) and throughout the whole brain (global). As random effects intercept and slope of RSFC for each individual were included in the models. Models were corrected for baseline age and sex, for longitudinal mean frame-displacement (FD) and for the interactions of those covariates with time. Unstandardized estimates (Est.), standard errors (SE), t-values and p-values (**$p<0.01$, *$p<0.05$) as well as marginal (R2m) and conditional (R2c) $R^2$ values are presented. Obs., observations; part, participants.

### eTable 7. Association between AD biomarkers and longitudinal changes in RSFC, using the Power parcellation

| Aβ | DMN | | | | SAL | | | | VAN | | | |
|---|---|---|---|---|---|---|---|---|---|---|---|---|
| n of obs. = 340 | | | | | | | | | | | | |
| n of part. = 91 | Est. | SE | t | p | Est. | SE | t | p | Est. | SE | t | p |
| Aβ x time | 0.0007 | 0.002 | 0.346 | 0.731 | 0.0016 | 0.003 | 0.539 | 0.592 | -0.0044 | 0.004 | -1.148 | 0.252 |
| age x time | 0.0003 | 0.002 | 0.169 | 0.867 | -0.0021 | 0.003 | -0.715 | 0.478 | 0.0053 | 0.004 | 1.403 | 0.162 |
| sex x time | 0.0058 | 0.004 | 1.376 | 0.175 | 0.0018 | 0.006 | 0.278 | 0.782 | 0.0031 | 0.008 | 0.369 | 0.712 |
| mean FD x time | 0.0013 | 0.002 | 0.729 | 0.468 | 0.0025 | 0.003 | 0.924 | 0.358 | 0.0007 | 0.004 | 0.195 | 0.845 |
| Aβ | -0.0008 | 0.003 | -0.275 | 0.784 | -0.0009 | 0.005 | -0.187 | 0.852 | -0.0093 | 0.007 | -1.327 | 0.188 |
| age | -0.0017 | 0.003 | -0.527 | 0.600 | 0.0045 | 0.005 | 0.885 | 0.378 | -0.0037 | 0.007 | -0.520 | 0.604 |
| sex | -0.0019 | 0.007 | -0.260 | 0.796 | -0.0069 | 0.012 | -0.589 | 0.557 | -0.0330 | 0.016 | -2.043 | 0.044* |
| mean FD | 0.0016 | 0.002 | 0.661 | 0.510 | 0.0090 | 0.004 | 2.455 | 0.015* | 0.0015 | 0.005 | 0.278 | 0.781 |
| time | -0.0067 | 0.004 | -1.856 | 0.069. | -0.0047 | 0.006 | -0.856 | 0.396 | 0.0000 | 0.007 | 0.000 | 1.000 |
| R2m | 0.014 | | | | 0.043 | | | | 0.046 | | | |
| R2c | 0.501 | | | | 0.563 | | | | 0.489 | | | |

| Aβ | FPN | | | | LIM | | | | DAN | | | |
|---|---|---|---|---|---|---|---|---|---|---|---|---|
| n of obs. = 340 | | | | | | | | | | | | |
| n of part. = 91 | Est. | SE | t | p | Est. | SE | t | p | Est. | SE | t | p |
| Aβ x time | 0.0018 | 0.002 | 0.806 | 0.421 | -0.0022 | 0.004 | -0.500 | 0.618 | 0.0034 | 0.003 | 1.020 | 0.312 |
| age x time | -0.0015 | 0.002 | -0.684 | 0.495 | 0.0024 | 0.004 | 0.546 | 0.587 | -0.0018 | 0.003 | -0.544 | 0.588 |
| sex x time | -0.0022 | 0.005 | -0.444 | 0.657 | 0.0035 | 0.010 | 0.358 | 0.721 | 0.0102 | 0.007 | 1.438 | 0.156 |
| mean FD x time | 0.0031 | 0.002 | 1.406 | 0.161 | 0.0039 | 0.004 | 0.903 | 0.369 | 0.0059 | 0.003 | 1.843 | 0.069. |
| Aβ | -0.0030 | 0.005 | -0.558 | 0.578 | 0.0077 | 0.009 | 0.884 | 0.379 | -0.0064 | 0.007 | -0.944 | 0.348 |
| age | 0.0047 | 0.005 | 0.866 | 0.389 | -0.0101 | 0.009 | -1.132 | 0.261 | -0.0132 | 0.007 | -1.902 | 0.060. |
| sex | 0.0012 | 0.012 | 0.096 | 0.924 | 0.0329 | 0.020 | 1.633 | 0.106 | -0.0309 | 0.016 | -1.971 | 0.052. |
| mean FD | 0.0043 | 0.003 | 1.239 | 0.216 | 0.0015 | 0.006 | 0.242 | 0.809 | 0.0077 | 0.005 | 1.603 | 0.110 |
| time | -0.0025 | 0.004 | -0.591 | 0.555 | -0.0043 | 0.008 | -0.518 | 0.606 | -0.0062 | 0.006 | -1.009 | 0.317 |
| R2m | 0.023 | | | | 0.035 | | | | 0.072 | | | |
| R2c | 0.631 | | | | 0.542 | | | | 0.587 | | | |

| Aβ | VIS | | | | SM | | | | global | | | |
|---|---|---|---|---|---|---|---|---|---|---|---|---|
| n of obs. = 340 | | | | | | | | | | | | |
| n of part. = 91 | Est. | SE | t | p | Est. | SE | t | p | Est. | SE | t | p |
| Aβ x time | -0.0012 | 0.005 | -0.262 | 0.794 | -0.0036 | 0.005 | -0.787 | 0.432 | 0.0010 | 0.006 | 0.172 | 0.864 |
| age x time | -0.0019 | 0.005 | -0.408 | 0.684 | 0.0030 | 0.005 | 0.657 | 0.512 | -0.0005 | 0.002 | -0.319 | 0.750 |
| sex x time | -0.0060 | 0.010 | -0.595 | 0.552 | 0.0125 | 0.010 | 1.254 | 0.211 | 0.0017 | 0.004 | 0.471 | 0.638 |
| mean FD x time | 0.0045 | 0.005 | 0.995 | 0.321 | 0.0015 | 0.004 | 0.331 | 0.741 | 0.0022 | 0.002 | 1.322 | 0.188 |
| Aβ | -0.0105 | 0.009 | -1.143 | 0.256 | -0.0062 | 0.009 | -0.675 | 0.501 | -0.0076 | 0.010 | -0.732 | 0.466 |
| age | -0.0039 | 0.009 | -0.417 | 0.678 | 0.0072 | 0.009 | 0.772 | 0.442 | -0.0005 | 0.003 | -0.149 | 0.882 |
| sex | -0.0221 | 0.021 | -1.050 | 0.297 | -0.0443 | 0.021 | -2.094 | 0.039* | -0.0115 | 0.007 | -1.585 | 0.117 |
| mean FD | 0.0076 | 0.007 | 1.140 | 0.255 | 0.0086 | 0.007 | 1.300 | 0.195 | 0.0048 | 0.002 | 2.087 | 0.038* |
| time | 0.0059 | 0.009 | 0.672 | 0.502 | -0.0023 | 0.009 | -0.274 | 0.784 | -0.0017 | 0.008 | -0.215 | 0.830 |
| R2m | 0.025 | | | | 0.057 | | | | 0.041 | | | |
| R2c | 0.530 | | | | 0.553 | | | | 0.515 | | | |

| *Tau* | DMN | | | | SAL | | | | VAN | | | |
|---|---|---|---|---|---|---|---|---|---|---|---|---|
| n of obs. = 340 | | | | | | | | | | | | |
| n of part. = 91 | Est. | SE | t | p | Est. | SE | t | p | Est. | SE | t | p |
| tau x time | -0.0007 | 0.002 | -0.365 | 0.716 | -0.0043 | 0.003 | -1.468 | 0.147 | -0.0025 | 0.004 | -0.626 | 0.532 |
| age x time | 0.0007 | 0.002 | 0.347 | 0.730 | -0.0004 | 0.003 | -0.154 | 0.878 | 0.0051 | 0.004 | 1.308 | 0.192 |
| sex x time | 0.0058 | 0.004 | 1.368 | 0.177 | 0.0019 | 0.006 | 0.307 | 0.760 | 0.0035 | 0.008 | 0.417 | 0.677 |
| mean FD x time | 0.0013 | 0.002 | 0.718 | 0.474 | 0.0024 | 0.003 | 0.876 | 0.383 | 0.0009 | 0.004 | 0.227 | 0.821 |
| tau | 0.0011 | 0.003 | 0.344 | 0.732 | 0.0017 | 0.005 | 0.335 | 0.738 | -0.0035 | 0.007 | -0.485 | 0.629 |
| age | -0.0021 | 0.003 | -0.658 | 0.513 | 0.0039 | 0.005 | 0.740 | 0.461 | -0.0045 | 0.007 | -0.608 | 0.545 |
| sex | -0.0018 | 0.007 | -0.249 | 0.804 | -0.0067 | 0.012 | -0.575 | 0.567 | -0.0297 | 0.016 | -1.832 | 0.070. |
| mean FD | 0.0016 | 0.002 | 0.659 | 0.510 | 0.0088 | 0.004 | 2.388 | 0.018* | 0.0015 | 0.005 | 0.284 | 0.777 |
| time | -0.0068 | 0.004 | -1.856 | 0.069. | -0.0049 | 0.005 | -0.916 | 0.364 | -0.0002 | 0.007 | -0.023 | 0.982 |
| R2m | 0.014 | | | | 0.045 | | | | 0.031 | | | |
| R2c | 0.502 | | | | 0.563 | | | | 0.486 | | | |

| *Tau* | FPN | | | | LIM | | | | DAN | | | |
|---|---|---|---|---|---|---|---|---|---|---|---|---|
| n of obs. = 340 | | | | | | | | | | | | |
| n of part. = 91 | Est. | SE | t | p | Est. | SE | t | p | Est. | SE | t | p |
| tau x time | -0.0003 | 0.002 | -0.117 | 0.907 | 0.0008 | 0.005 | 0.173 | 0.863 | -0.0003 | 0.003 | -0.095 | 0.925 |
| age x time | -0.0010 | 0.002 | -0.451 | 0.652 | 0.0016 | 0.004 | 0.366 | 0.715 | -0.0009 | 0.003 | -0.266 | 0.791 |
| sex x time | -0.0021 | 0.005 | -0.431 | 0.667 | 0.0028 | 0.010 | 0.292 | 0.771 | 0.0107 | 0.007 | 1.496 | 0.140 |
| mean FD x time | 0.0031 | 0.002 | 1.389 | 0.166 | 0.0039 | 0.004 | 0.902 | 0.370 | 0.0059 | 0.003 | 1.821 | 0.072. |
| tau | 0.0006 | 0.005 | 0.114 | 0.909 | 0.0096 | 0.009 | 1.082 | 0.282 | -0.0052 | 0.007 | -0.749 | 0.456 |
| age | 0.0040 | 0.006 | 0.717 | 0.475 | -0.0111 | 0.009 | -1.225 | 0.224 | -0.0131 | 0.007 | -1.856 | 0.067. |
| sex | 0.0016 | 0.012 | 0.129 | 0.898 | 0.0307 | 0.020 | 1.533 | 0.129 | -0.0295 | 0.016 | -1.884 | 0.063. |
| mean FD | 0.0043 | 0.003 | 1.237 | 0.217 | 0.0014 | 0.006 | 0.223 | 0.824 | 0.0078 | 0.005 | 1.619 | 0.106 |
| time | -0.0026 | 0.004 | -0.620 | 0.536 | -0.0037 | 0.008 | -0.446 | 0.657 | -0.0067 | 0.006 | -1.088 | 0.282 |
| R2m | 0.020 | | | | 0.036 | | | | 0.066 | | | |
| R2c | 0.633 | | | | 0.539 | | | | 0.584 | | | |

| *Tau* | VIS | | | | SM | | | | global | | | |
|---|---|---|---|---|---|---|---|---|---|---|---|---|
| n of obs. = 340 | | | | | | | | | | | | |
| n of part. = 91 | Est. | SE | t | p | Est. | SE | t | p | Est. | SE | t | p |
| tau x time | -0.0013 | 0.005 | -0.263 | 0.793 | -0.0029 | 0.005 | -0.605 | 0.546 | 0.0000 | 0.002 | 0.024 | 0.981 |
| age x time | -0.0018 | 0.005 | -0.383 | 0.702 | 0.0029 | 0.005 | 0.632 | 0.528 | -0.0005 | 0.002 | -0.280 | 0.780 |
| sex x time | -0.0055 | 0.010 | -0.544 | 0.587 | 0.0131 | 0.010 | 1.316 | 0.189 | 0.0018 | 0.004 | 0.502 | 0.616 |
| mean FD x time | 0.0046 | 0.005 | 1.011 | 0.313 | 0.0016 | 0.004 | 0.357 | 0.722 | 0.0022 | 0.002 | 1.327 | 0.186 |
| tau | -0.0089 | 0.009 | -0.956 | 0.342 | -0.0152 | 0.009 | -1.644 | 0.104 | -0.0020 | 0.003 | -0.632 | 0.529 |
| age | -0.0035 | 0.010 | -0.368 | 0.713 | 0.0100 | 0.009 | 1.068 | 0.288 | -0.0004 | 0.003 | -0.118 | 0.906 |
| sex | -0.0186 | 0.021 | -0.885 | 0.379 | -0.0406 | 0.021 | -1.950 | 0.054. | -0.0108 | 0.007 | -1.498 | 0.138 |
| mean FD | 0.0077 | 0.007 | 1.158 | 0.248 | 0.0086 | 0.007 | 1.309 | 0.192 | 0.0049 | 0.002 | 2.101 | 0.037* |
| time | 0.0055 | 0.009 | 0.632 | 0.528 | -0.0027 | 0.009 | -0.321 | 0.748 | -0.0005 | 0.003 | -0.169 | 0.866 |
| R2m | 0.021 | | | | 0.068 | | | | 0.038 | | | |
| R2c | 0.530 | | | | 0.550 | | | | 0.515 | | | |

Using linear mixed effects models, we test whether global amyloid-β (Aβ) and entorhinal *tau* deposition were associated with a change in resting-state functional connectivity (RSFC) within different functional networks that were defined by the Power parcellation atlas (DMN, default mode network; SAL, salience network; VAN, ventral attention network; FPN, fronto-parietal network; LIM, limbic network; VIS, visual network and SM, somatomotor network) and throughout the whole brain (global). As random effects intercept and slope of RSFC for each individual were included in the models. Models were corrected for baseline age and sex, for longitudinal mean frame-displacement (FD) and for the interactions of those covariates with time. Unstandardized estimates (Est.), standard errors (SE), t-values and p-values (*p<0.05) as well as marginal (R2m) and conditional (R2c) $R^2$ values are presented. Obs., observations; part, participants.
